# Cell Surface Nucleocapsid Protein Expression: A Betacoronavirus Immunomodulatory Strategy

**DOI:** 10.1101/2023.02.24.529952

**Authors:** Alberto Domingo López-Muñoz, Jefferson J.S. Santos, Jonathan W. Yewdell

## Abstract

We recently reported that SARS-CoV-2 Nucleocapsid (N) protein is abundantly expressed on the surface of both infected and neighboring uninfected cells, where it enables activation of Fc receptor-bearing immune cells with anti-N antibodies (Abs) and inhibits leukocyte chemotaxis by binding chemokines (CHKs). Here, we extend these findings to N from the seasonal human coronavirus (HCoV)-OC43, which is also robustly expressed on the surface of infected and non-infected cells by binding heparan-sulfate/heparin (HS/H). HCoV-OC43 N binds with high affinity to the same set of 11 human CHKs as SARS-CoV-2 N, but also to a non-overlapping set of 6 cytokines (CKs). As with SARS-CoV-2 N, HCoV-OC43 N inhibits CXCL12β-mediated leukocyte migration in chemotaxis assays, as do all highly pathogenic and endemic HCoV N proteins. Together, our findings indicate that cell surface HCoV N plays important evolutionary conserved roles in manipulating host innate immunity and as a target for adaptive immunity.

## INTRODUCTION

Over just twenty years three highly pathogenic HCoVs have emerged with the latest, SARS-CoV-2, causing millions of deaths and economic havoc. Given the high potential of newly emerging HCoVs to be introduced from the enormous animal reservoir, it is critical to deepen the knowledge of HCoVs life cycle and immune evasion strategies.

The four common seasonal HCoVs (OC43, HKU1, NL63 and 229E) cause typically mild upper respiratory infections. As a betacoronavirus, SARS-CoV-2 is more closely related to HCoV-OC43 and HKU1 than to the alphacoronaviruses 229E and NL63 (*1*). For all HCoVs, viral entry is mediated by the Spike (S) receptor-fusion glycoprotein. NL63, SARS-CoV-2 and SARS-CoV-1 S proteins use angiotensin-converting enzyme 2 (ACE2) to bind cells, while MERS-CoV binds dipeptidyl peptidase 4 (DPP4), and 229E uses aminopeptidase N (CD13) (*2*). Specific protein receptors have not been identified yet for HCoV-OC43 and HKU1, whose S protein bind terminal 9-O-acetylsialic acid for entry (*3*).

HCoV N binds viral RNA and plays pivotal roles in packaging and transcribing viral RNA (*4*). While the sequence identity between SARS-CoV2 and HCoV-OC43 N is the lowest (38%) among HCoVs, domain architecture and overall structure is highly conserved across them (*5, 6*). N from all animal and HCoV strains studied suppress interferon (IFN) responses using multiples strategies (*7–10*).

The cell surface expression of viral RNA- and DNA-binding proteins dates to initial polyclonal Ab detection of retrovirus *gag* (*11*) and polyoma virus T antigen in the 1970s (*12*). These findings were extended to influenza virus N using polyclonal Abs (*13*), and definitively established with monoclonal Abs (mAbs) (*14*). Similar findings were made using mAbs specific for surface N expressed by vesicular stomatitis (*15*), lymphocytic choriomeningitis (*16*), human immunodeficiency (*17*), respiratory syncytial (*18*), and measles viruses (*19*). For the latter two viruses, respectively, cell surface N has been reported to impair the immunological synapse formation with T cells and bloc IL-12 secretion (*18–20*).

Recently, our group reported that N synthesized from cells infected with SARS-CoV-2 or transfected with a N encoding plasmid is robustly released from cells, binding to infected and non-infected neighboring cells, where it modulates innate and adaptive immunity (*21*). It was previously reported that mouse hepatitis coronavirus N is present on the cell surface where it enables complement-mediated lysis of infected cells and enables protection against disease (*22, 23*). Here, we extend these findings to the seasonal HCoV-OC43 N protein.

## RESULTS

### Surface HCoV-OC43 N is consistently expressed across infected cell lines and human airway epithelium (HAE)

We studied cell surface expression of HCoV-OC43 N by imaging HEK-293FT, MRC-5 and Vero cells 24 h post-infection (hpi). To exclusively detect extracellular N, we incubated live cells with primary and fluorescent secondary Abs at 4°C prior to fixation and mounting for confocal imaging. We observed a clear surface N staining over mock-infected background levels in the three cell types examined, as well as for the S protein (Fig. 1A, maximum intensity projection images of *z*-stack). We noted a remarkable degree of colocalization between N and S in Vero cells, as we previously reported for SARS-CoV-2-infected Vero cells (*21*).

**Fig. 1.**
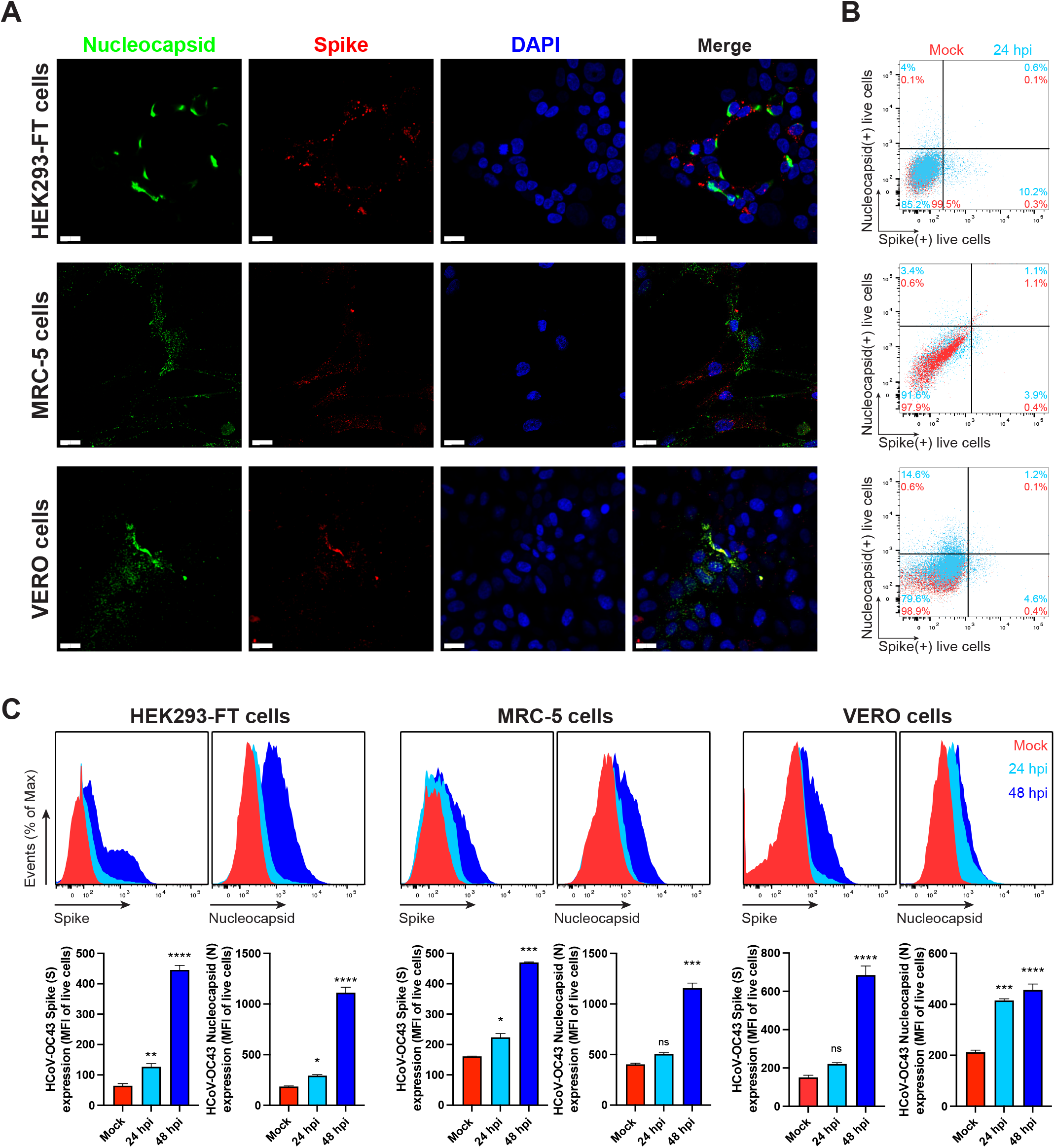
N is expressed on the surface of live HCoV-OC43 infected cells. **(A)** Maximum intensity projections of laser confocal microscopy z-stack images of infected HEK-293FT, MRC-5 and Vero cells with HCoV-OC43, stained live at 24 hpi (MOI = 1). Scale bar = 20 μm. Images are representative of at least three independent experiments with similar results. **(B)** Flow cytometry analyses of viable infected cells (MOI = 1), stained live at 24 hpi for HCoV-OC43 S and N proteins. Representative dot plots of flow cytometry analyses showing double staining of surface S and N, indicating the percentage of the gated cell population for each quadrant of the double staining. Data are representative of at least three independent experiments, each performed with triplicate samples. **(C, D)** Time course of surface S and N proteins expression in live infected cells with HCoV-OC43 at 24 and 48 hpi (MOI = 1). For each infected cell type, the following is shown: histogram overlays of surface S and N proteins, as well as the mean fluorescent intensity (MFI) is plotted showing mean +/- SEM (n = 3). One-way ANOVA and Dunnett s Multiple comparison test were used to compare infected conditions against mock-infected cells: ns (nonsignificant) *p* > 0.05, * *p* < 0.05, ** *p* < 0.01, *** *p* < 0.001, **** *p* < 0.0001. Data are representative of one experiment out of at least three independent experiments performed in triplicate.

For a more quantitative measurement of N surface expression, we performed flow cytometry using live cells 24 hpi with HCoV-OC43. We detected cell surface N in HEK-293FT, MRC-5, Vero (Fig. 1B, 1C), RD and BHK-21 cells (Fig. S1). Surface N was more robustly detected in HEK-293FT, MRC-5, Vero and CHO-K1 cells 48 hpi (Fig. 1C, S1). Remarkably, depending on the cell type, we noticed an increment in the fraction of cells expressing N but not S over time (Fig. S2). This fraction of N+/S-cells was higher that the fraction of N+/S+ cells in all cell types examined but RD cells (Fig. 1B, S1), ranging from 1% to 14.6% at 24 hpi. This is consistent with our previous findings in SARS-CoV-2 infected cells, and most likely due to the transfer of N released from infected to non-infected cells.

To determine whether N cell surface expression occurs in HAE, we infected human nasal (MucilAir™) and bronchial (SmallAir™) airway epithelial cells cultured at the air-liquid interface with HCoV-OC43 or SARS-CoV-2. Via flow cytometry, we detected a discrete but significant surface signal for N and S proteins in HCoV-OC43-infected MucilAir™ and SmallAir™ cultures 72 hpi (Fig. 2). SARS-CoV-2-infected MucilAir™ cultures also showed significant surface signal for N and S proteins 72 hpi (Fig. S3).

**Fig. 2.**
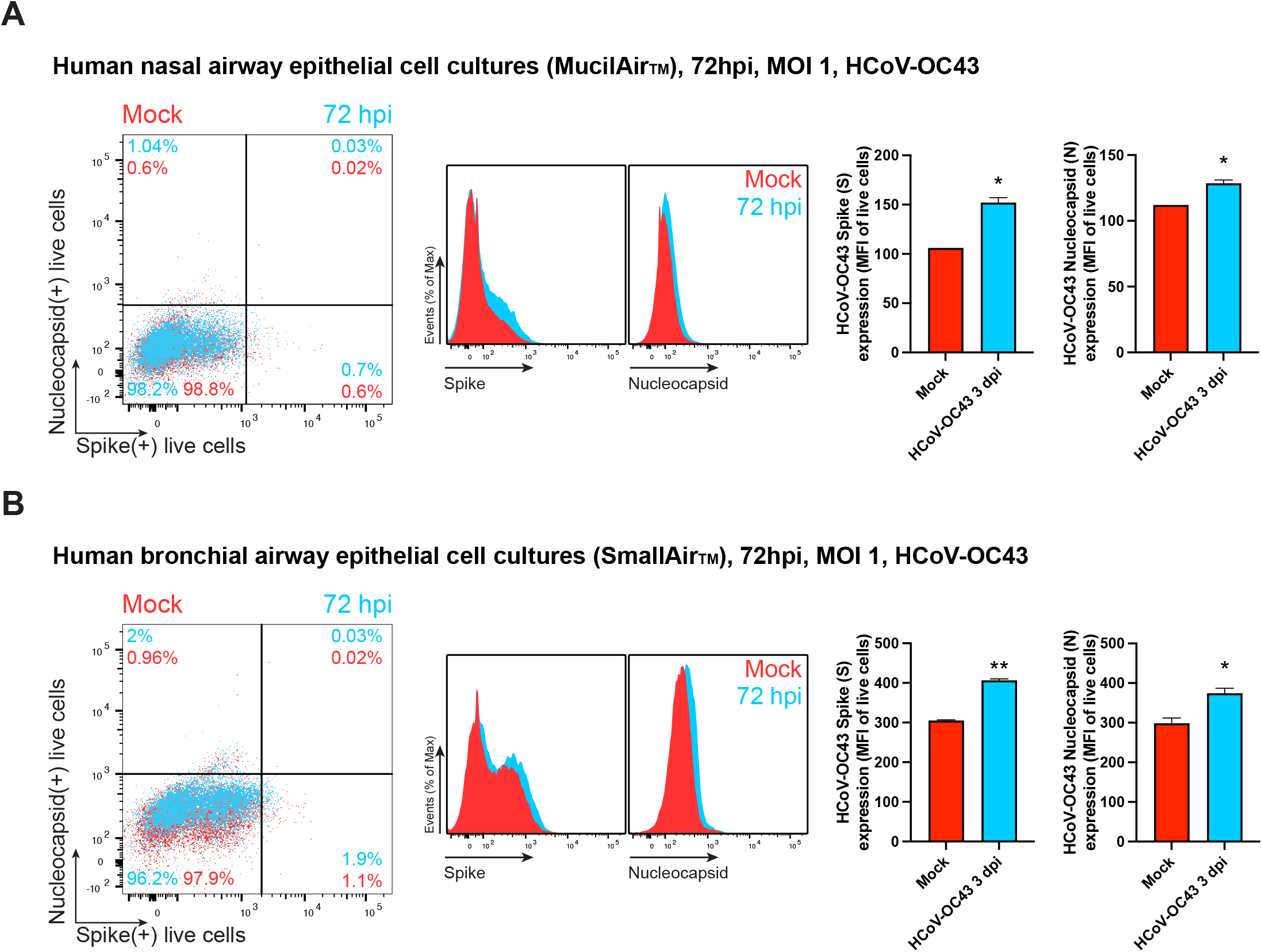
N is expressed on the surface of live HCoV-OC43 infected HAE. Flow cytometry analyses of human nasal **(A)** and bronchial **(B)** airway epithelial cells infected with HCoV-OC43 (MOI = 1), stained live at 72 hpi to detect cell surface S and N protein. Representative dot plots of flow cytometry analyses showing double staining of surface S and N, indicating the percentage of the gated cell population for each quadrant. For each infection, the following is shown: histogram overlays of surface S and N proteins, as well as the MFI is plotted showing mean +/- SEM (n = 2). * *p* < 0.05, ** *p* < 0.01 by Student s two-tailed unpaired *t*-test. Data are representative of one experiment out of two independent experiments performed in duplicate.

Together, these and previous results indicate that N is robustly localized on the surface of cells infected with human betacoronaviruses.

### Electrostatic association with surface HS/H mediates HCoV-OC43 N binding independently of other viral genes

We examined the surface expression of N in synthesizing cells following transient transfection with an expression plasmid. Live HEK-293FT, BHK-21 and CHO-K1 transfected cells showed significant N Ab binding in flow cytometry over background levels from cells transfected with a control plasmid expressing eGFP (Fig. 3A, S4A). Staining with different N Abs showed similar and consistent results across independent experiments, supporting the specificity of staining for N.

**Fig. 3.**
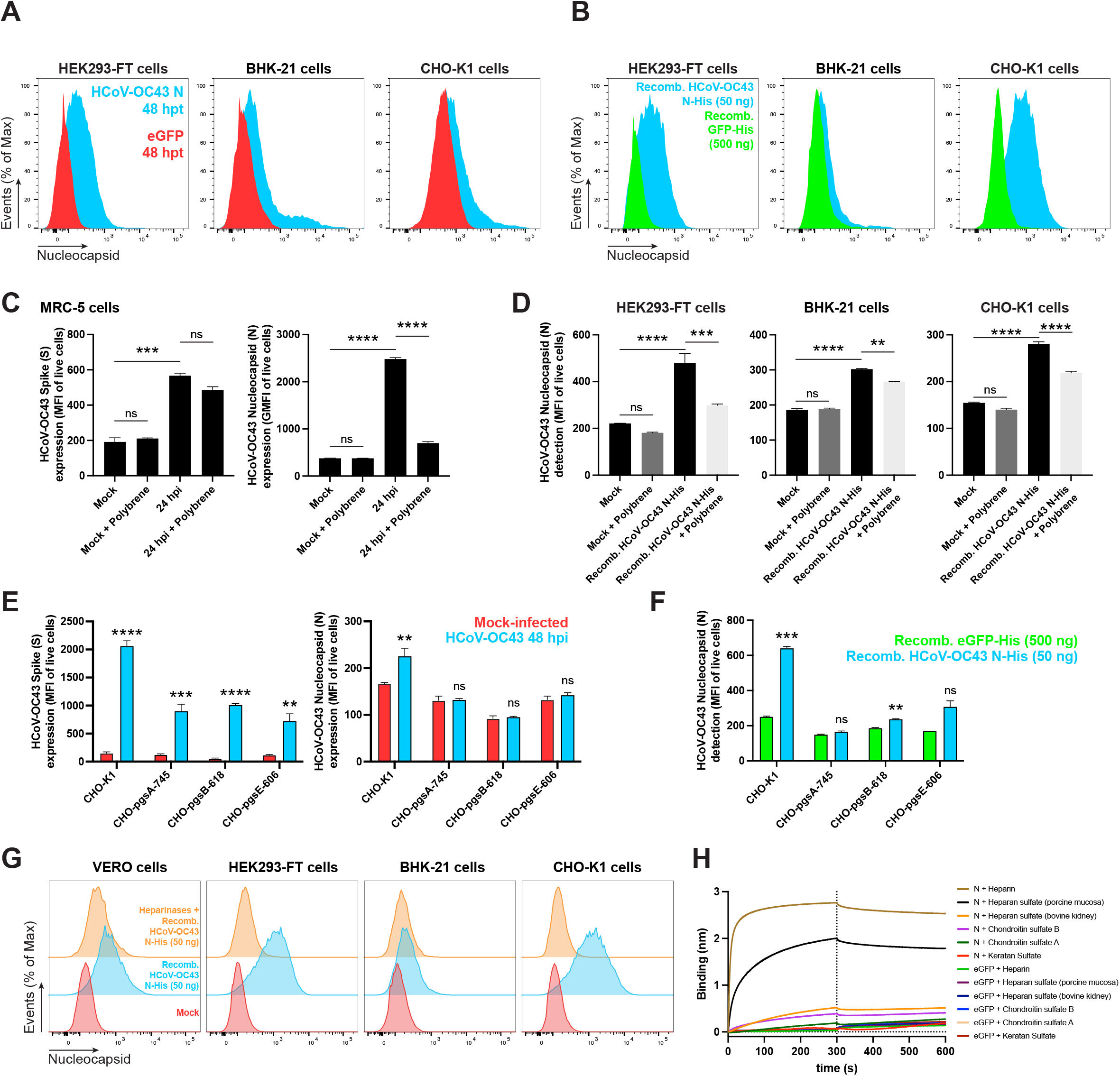
Cell surface binding of HCoV-OC43 N occurs independently of other viral genes and is mediated by electrostatic interactions with HS/H. **(A)** Histogram overlays of surface N protein expression of live HEK293-FT, BHK-21 and CHO-K1 cells transiently transfected with a plasmid encoding eGFP (negative control) or N protein, detected with Abs by flow cytometry. **(B)** Histogram overlays of exogenous rN binding to HEK293-FT, BHK-21 and CHO-K1 cells, incubated with recombinant eGFP (negative control) or N protein for 15 min, washed twice, stained live with Abs, and analyzed by flow cytometry. **(C)** Electric charge neutralization assay with a cationic polymer (polybrene) on infected cells. MRC-5 cells were infected with HCoV-OC43 (MOI = 10), washed twice, incubated with 10 μg/ml of polybrene, washed twice, stained live with Abs, and analyzed by flow at 24 hpi. **(D)** Electric charge neutralization assays with exogenous rN. HEK293-FT, BHK-21 and CHO-K1 cells were incubated with 50 ng of rN protein for 15 min, washed twice, incubated with 10 μg/ml of polybrene, washed twice, stained live with Abs and analyzed by flow. **(E)** GAG-deficient CHO cells were infected with HCoV-OC43 (MOI = 1), washed twice, incubated with 10 μg/ml of polybrene, washed twice, stained live with Abs, and analyzed by flow at 48 hpi. **(F)** GAG-deficient CHO cells were incubated with recombinant eGFP or rN protein for 15 min, washed twice, stained live with Abs, and analyzed by flow cytometry. **(G)** Heparinase treatment significantly abrogates the cell ability to bind and retain the N protein. Flow cytometry histogram semi-overlays of Vero, HEK293-FT, BHK-21 and CHO-K1 cells treated with heparinases for 1 h, washed twice, incubated with 50 ng of rN protein for 15 min, washed twice, stained live with Abs, and analyzed. **(H)** BLI sensorgrams from binding assays of sulfated GAGs to immobilized N or eGFP proteins. Streptavidin-coated biosensors were loaded with equivalent amounts of N or eGFP, measuring their ability to bind each GAG. Sensorgrams show association and dissociation phases, where the vertical dotted line indicates the end of the association step. In **(C, D, E, F)** the MFI of detected surface N protein from live cells is plotted, showing mean +/- SEM (n = 2). For **(C, D)** One-way ANOVA and Dunnett s Multiple comparison test were used to compare all conditions against mock cells: ns (nonsignificant) *p* > 0.05, ** *p* < 0.01, *** *p* < 0.001, **** *p* < 0.0001. In **(E, F)**, ns (nonsignificant statistically) *p* > 0.01, ** *p* < 0.01, *** *p* < 0.001, **** *p* < 0.0001 by Student s two-tailed unpaired *t*-test. The analyses were repeated with different protein preparations, and one representative assay out of at least three independent assays performed in duplicate is shown.

We next incubated HEK-293FT, BHK-21 and CHO-K1 cells with purified recombinant N (rN) for 15 min at 37 C°. Following staining with anti-N Abs, flow cytometry assays revealed strong surface staining with anti-N Abs compared to control cells incubated with GFP as a control protein (Fig. 3B, S4B). The degree of staining varied between cell lines, reaching up to a 10-fold increase over background levels for CHO-K1 cells.

The RNA-binding domains of the N protein are heavily positively charged, whose electrostatic association with the negatively charged viral RNA contributes to the free energy of the interaction (*24, 25*). The negative charge of the cellular surface results mainly from glycosaminoglycans (GAGs). Among these, H, a highly sulfated form of HS, has the highest negative charge of known biomolecules (*26*). We reported charge-based SARS-CoV-2 N binding to the cell surface by using polybrene, a cationic polymer that neutralizes surface electrostatic charges. Treating HCoV-OC43-infected MRC-5 cells with polybrene removed cell surface N detected by flow cytometry 24 hpi. Importantly, the amount of S, a membrane anchored protein, was not diminished (Fig. 3C, S4C). Similarly, polybrene removed a substantial fraction of exogenous rN bound to the cell surface of live HEK-293FT, BHK-21 and CHO-K1 cells (Fig. 3D, S4D).

We previously showed that GAGs, specifically HS/H, mediate SARS-CoV-2 N cell surface binding. We used a panel of GAG-deficient CHO cells (*27*) to determine the contribution of GAGs to N cell surface binding. CHO-pgsA-745 cells do not express xylosyltransferase (*28*), while CHO-pgsB-618 cells are deficient in β-1,4-galactosyltransferase 7 activity (*29*), resulting in in the complete absence of GAGs in either cell type. CHO-pgsE-606 cells are partially deficient synthesizing HS but can synthesize other GAGs (*30, 31*). We performed flow cytometry analyses of live infected wild-type (CHO-K1) and GAG-deficient CHO cells 48 hpi with HCoV-OC43. While surface S was significantly detected on CHO-K1 and every GAG-deficient CHO cell, N was only detected on the surface of CHO-K1 cells (Fig. 3E, S4E). Each of the GAG-deficient CHO cells also failed to bind and retain exogenous rN over background levels with recombinant GFP (Fig. 3F, S4F). Treating cells with heparinases (I, II, and III combined) nearly completely depolymerizes surface HS/H polysaccharide chains to soluble disaccharides. Heparinases treatment of Vero, HEK293-FT, BHK-21 and CHO-K1 cells significantly reduced binding and retention of exogenous rN by flow cytometry (Fig. 3G, S4G).

The most ubiquitous sulfated GAGs on the cell surface and in the extracellular matrix are HS/H, and chondroitin sulfate A/B (*32*). By biolayer interferometry (BLI), we confirmed specific N binding to HS/H, demonstrating N nanomolar affinity exclusively to these complex highly sulfated GAGs (Fig. 3H, S5, Table 1).

**Table 1.** Kinetic analysis of HCoV-OC43 N protein binding to HS and H by BLI.

| GAG | $K_D$ (M) | $k_{on}$ (1/M.s) | $K_{off}$ (1/s) | $R^2$ |
| --- | --- | --- | --- | --- |
| Heparan sulfate (porcine mucosa) | $1.5 \times 10^{-8} \pm 1.6 \times 10^{-10}$ | $4.2 \times 10^4 \pm 1.9 \times 10^2$ | $6.2 \times 10^{-4} \pm 5.9 \times 10^{-6}$ | 0.996 |
| Heparin | $8.9 \times 10^{-10} \pm 1.3 \times 10^{-11}$ | $2.9 \times 10^5 \pm 7 \times 10^2$ | $2.6 \times 10^{-4} \pm 3.8 \times 10^{-6}$ | 0.998 |

Values reported represent the average of the global fit ( ± error) to the data from at least three independent BLI kinetic assays performed with different N protein preparations. Kinetics parameters were determined by using the heterogeneous ligand (2:1) binding model.

Together, these results show that surface electrostatic charges given by GAGs, particularly by HS/H, play a fundamental role mediating betacoronavirus N protein binding to the cell surface.

### HCoV-OC43 N is trans-expressed on the cell surface of non-expressing cells

The increasing number of cells expressing N but not S over time after infection in HCoV-OC43 immunofluorescence and flow cytometry experiments (Figs. 1, 2, S1-S3), is consistent with transfer of N from infected to uninfected cells. To examine whether HCoV-OC43 N can be transferred from expressing to neighboring non-expressing cells, we co-cultured transiently N-transfected (donor) and non-transfected (recipient) cells at a ratio of 1 to 9 (donor to recipient). Pre-straining non-transfected cells with CellTrace™ Violet enabled their flow identification after co-culture (Fig. S6). Consistent with our previous findings for SARS-CoV-2 N (*21*), co-cultured non-transfected recipient cells exhibited significant amounts of surface N after overnight incubation with donor cells. Remarkably, for HEK293-FT cells, recipient cells had higher levels of cell surface N than donor cells (Fig. 4).

**Fig. 4.**
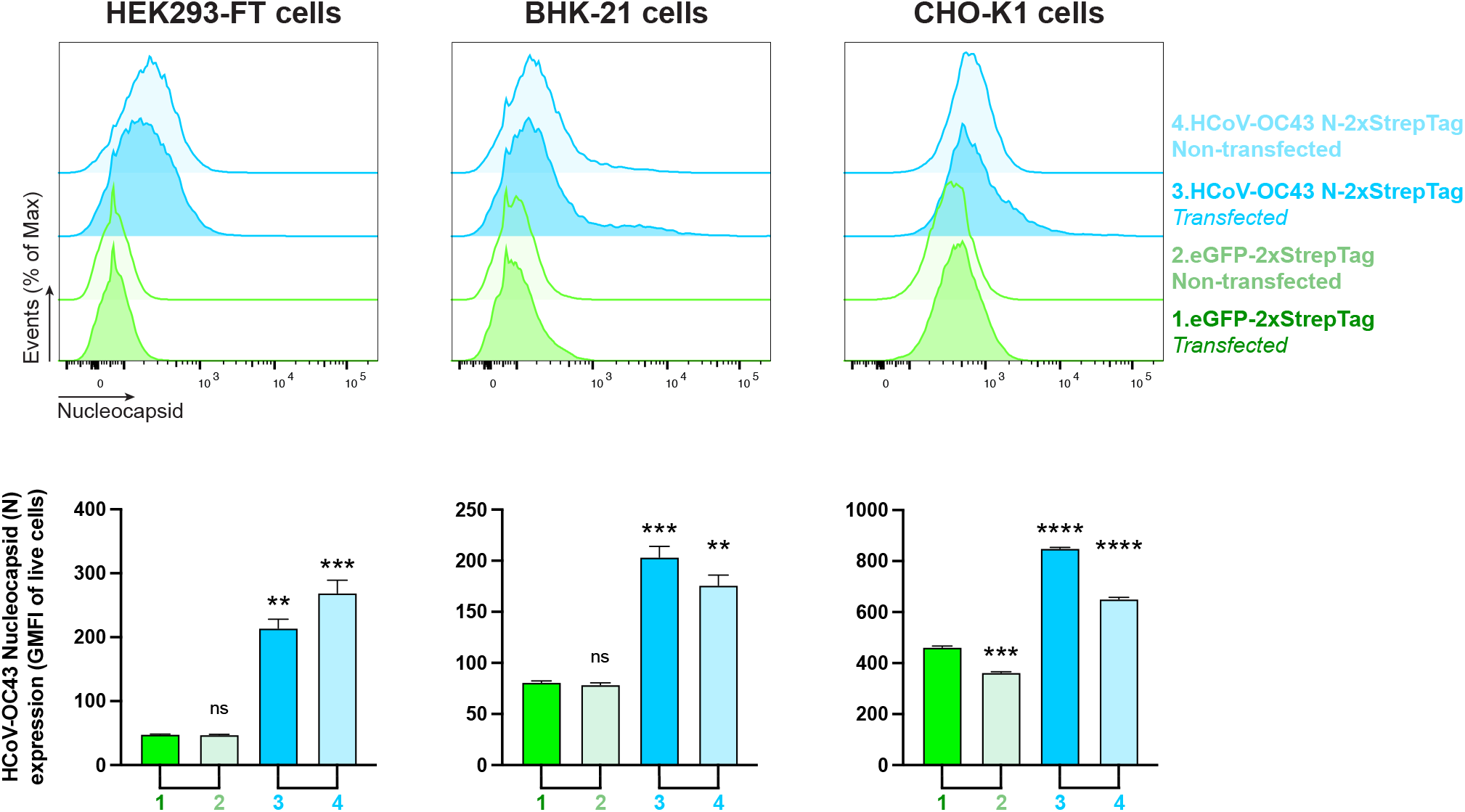
Cell surface HCoV-OC43 N intercellular transfer is independent of infection. Flow cytometry analyses of N transfer assays between transfected (donor) and non-transfected (recipient) co-cultured cells. Cells were transiently transfected with a plasmid encoding eGFP or the N protein. After 24 h, non-transfected cells were stained with CellTraceTM Violet prior to be co-cultured with their transfected counterparts. Cells were stained live after 12 h with Abs and analyzed. For each assay, the following is shown: histogram semi-overlays of surface N protein of live cells, as well as the MFI is plotted showing mean +/- SEM (n = 3). One representative experiment of at least three independent experiments performed in triplicate is shown. One-way ANOVA and Dunnett s Multiple comparison test were used to compare all conditions against eGFP-transfected cells: ns (nonsignificant) *p* > 0.05, ** *p* < 0.01, *** *p* < 0.001, **** *p* < 0.0001.

Together, these and previous findings indicate that N biosynthesis leads to its robust transfer to non-synthesizing cells.

### HCoV-OC43 N inhibits CHK-mediated leukocyte migration

Does HCoV-OC43 N interfere with CHK signaling as observed with the SARS-CoV-2 N protein? Using BLI, we assessed the binding of immobilized HCoV-OC43 rN to 64 human CKs. HCoV-OC43 N bound with high affinity to the same set of 11 human CHKs as SARS-CoV-2 N: CCL5, CCL11, CCL21, CCL26, CCL28, CXCL4, CXCL9, CXCL10, CXCL11, CXCL12β, and CXCL14. HCoV-OC43 N also bound to 6 additional CKs: CCL13, CCL20, CCL25, CXCL12α, CXCL13 and IL27 (Fig. 5A). N bound human CKs with high affinity, ranging from micromolar to nanomolar affinities (Table 2). Similarly to what we reported for SARS-CoV-2 N, kinetic curves of HCoV-OC43 N binding to each CK were also biphasic, deviating from first-order binding (1:1) and showing binding heterogeneity (Fig. S7) (*33*). None of the other CKs tested in the panel interacted with N with affinities higher than that observed for immobilized eGFP (Fig. S8A).

**Fig. 5.**
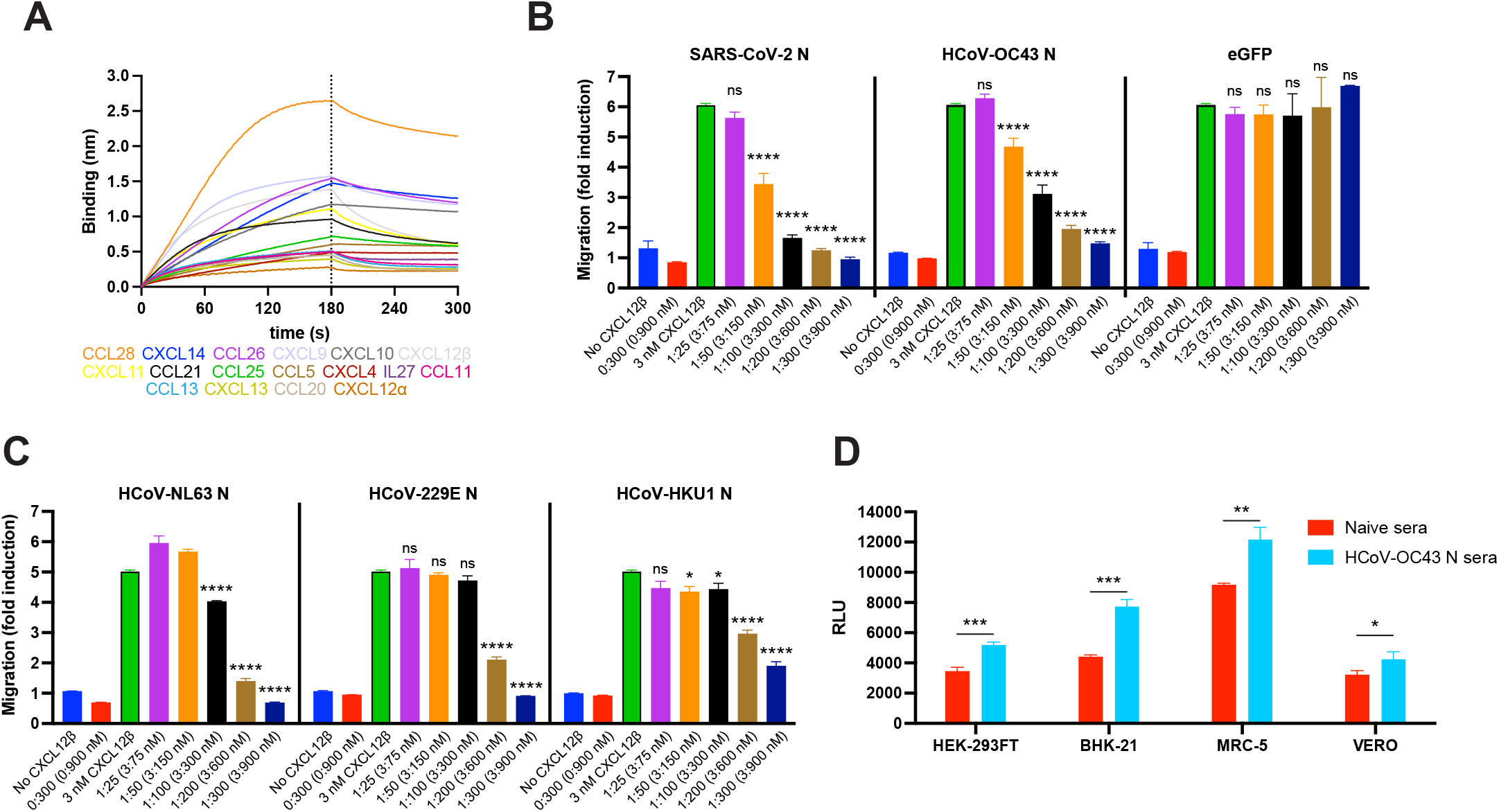
HCoV-OC43 N protein inhibits CHK function and is target for Ab Fc-based immunity. **(A, B)** N binds human CKs with high affinity and inhibits in vitro CHK-mediated leukocyte migration. **(A)** BLI sensorgrams of binding assays showing association and dissociation phases of the interaction between N protein and 17 positively bound CKs at a concentration of 100 nM out of 64 human cytokines tested. The dotted line indicates the end of the association step. The analyses were repeated with different purified rN protein preparations. One representative assay of three independent assays is shown. **(B)** HCOV-OC43 N blocks CXCL12β chemotaxis of MonoMac-1 cells, similarly to SARS-CoV-2 N. **(C)** N proteins from other endemic HCoVs also block CXCL12β chemotaxis of MonoMac-1 cells. In **(B, C)** CXCL12β was incubated alone (migration baseline, green bars) or in the presence of the indicated viral protein, in the lower chamber of transwell migration devices. Migrated cells from the top chamber were detected in the lower chamber at the end of the experiment. The induction of migration shows mean +/- SEM (n = 3) from one representative assay performed in triplicate out of at least three independent assays. One-way ANOVA and Dunnett s Multiple comparison test were used to compare all conditions (except no CHK and viral protein alone conditions) against migration induced by CHK alone (green bars): ns (nonsignificant) *p* > 0.05, * *p* < 0.05, ** *p* < 0.01, *** *p* < 0.001, **** *p* < 0.0001. **(D)** Cell surface HCoV-OC43 N protein is a target for Ab-based immunity. ADCC reporter bioassays were performed on HCoV-OC43-infected HEK-293FT, BHK-21, MRC-5 and Vero cells (24 hpi, MOI =1) using pooled sera from five mice immunized with HCoV-OC43 rN or from five naïve mice, and Jurkat effector-reporter cells. After overnight incubation, luciferase expression to gauge cell activation was measured. Data show mean +/- SEM (n = 3) of one representative assay out of three independent experiments performed in triplicate. * *p* < 0.05, ** *p* < 0.01, *** *p* < 0.001 by Student s two-tailed unpaired *t*-test.

**Table 2.** Kinetic analysis of HCoV-OC43 N protein binding to human CKs by BLI.

| CKs | K<sub>D</sub> (M) | k<sub>on</sub> (1/M.s) | K<sub>off</sub> (1/s) | R<sup>2</sup> |
| --- | --- | --- | --- | --- |
| CCL5 | 5.6x10<sup>-5</sup> ± 3.5x10<sup>-5</sup> | 2.1x10<sup>3</sup> ± 2.8x10<sup>2</sup> | 1.5x10<sup>-2</sup> ± 3.4x10<sup>-3</sup> | 0.989 |
| CCL11 | 1.4x10<sup>-6</sup> ± 4.8x10<sup>-8</sup> | 1.7x10<sup>4</sup> ± 4.2x10<sup>2</sup> | 2.1x10<sup>-2</sup> ± 4.2x10<sup>-4</sup> | 0.979 |
| CCL13 | 1.6x10<sup>-6</sup> ± 5x10<sup>-8</sup> | 1.9x10<sup>4</sup> ± 4.2x10<sup>2</sup> | 2.5x10<sup>-2</sup> ± 4.9x10<sup>-4</sup> | 0.967 |
| CCL20 | 1.8x10<sup>-6</sup> ± 1.4x10<sup>-7</sup> | 1.8x10<sup>4</sup> ± 6.3x10<sup>2</sup> | 1.7x10<sup>-2</sup> ± 4.9x10<sup>-4</sup> | 0.953 |
| CCL21 | 2.9x10<sup>-7</sup> ± 4.8x10<sup>-9</sup> | 6.7x10<sup>4</sup> ± 1.3x10<sup>3</sup> | 1.2x10<sup>-2</sup> ± 2x10<sup>-4</sup> | 0.963 |
| CCL25 | 6.1x10<sup>-7</sup> ± 2.8x10<sup>-8</sup> | 8.3x10<sup>3</sup> ± 1.5x10<sup>2</sup> | 4.8x10<sup>-3</sup> ± 1.9x10<sup>-4</sup> | 0.993 |
| CCL26 | 6.2x10<sup>-7</sup> ± 2.6x10<sup>-8</sup> | 2x10<sup>4</sup> ± 1.9x10<sup>2</sup> | 1.4x10<sup>-2</sup> ± 5.5x10<sup>-4</sup> | 0.991 |
| CCL28 | 2.7x10<sup>-8</sup> ± 5.1x10<sup>-10</sup> | 6x10<sup>4</sup> ± 6.4x10<sup>2</sup> | 1.5x10<sup>-3</sup> ± 2x10<sup>-5</sup> | 0.956 |
| CXCL4 | 1.7x10<sup>-7</sup> ± 1.6x10<sup>-8</sup> | 5.3x10<sup>3</sup> ± 3.3x10<sup>2</sup> | 8.8x10<sup>-4</sup> ± 6.8x10<sup>-5</sup> | 0.993 |
| CXCL9 | 2.9x10<sup>-8</sup> ± 9.1x10<sup>-10</sup> | 8.2x10<sup>4</sup> ± 1.5x10<sup>3</sup> | 2.2x10<sup>-3</sup> ± 6.2x10<sup>-5</sup> | 0.96 |
| CXCL10 | 3.7x10<sup>-7</sup> ± 4.3x10<sup>-8</sup> | 1x10<sup>4</sup> ± 2x10<sup>2</sup> | 3.7x10<sup>-3</sup> ± 4.2x10<sup>-4</sup> | 0.997 |
| CXCL11 | 7.3x10<sup>-7</sup> ± 2.5x10<sup>-8</sup> | 1.8x10<sup>4</sup> ± 2.8x10<sup>2</sup> | 1.2x10<sup>-2</sup> ± 3.8x10<sup>-4</sup> | 0.98 |
| CXCL12α | 4.6x10<sup>-10</sup> ± 2.4x10<sup>-10</sup> | 6.2x10<sup>4</sup> ± 1.4x10<sup>3</sup> | 3.7x10<sup>-5</sup> ± 1.4x10<sup>-5</sup> | 0.95 |
| CXCL12β | 1.5x10<sup>-6</sup> ± 3.8x10<sup>-8</sup> | 2.4x10<sup>4</sup> ± 3.3x10<sup>2</sup> | 1.9x10<sup>-2</sup> ± 2.1x10<sup>-4</sup> | 0.963 |
| CXCL13 | 1.4x10<sup>-7</sup> ± 9.2x10<sup>-9</sup> | 4.4x10<sup>4</sup> ± 9.2x10<sup>2</sup> | 4.6x10<sup>-3</sup> ± 2.7x10<sup>-4</sup> | 0.902 |
| CXCL14 | 5.8x10<sup>-7</sup> ± 2.6x10<sup>-8</sup> | 1.2x10<sup>4</sup> ± 2.6x10<sup>2</sup> | 6.3x10<sup>-3</sup> ± 2.6x10<sup>-4</sup> | 0.988 |
| IL27 | 5.5x10<sup>-8</sup> ± 1x10<sup>-9</sup> | 5.1x10<sup>4</sup> ± 1.8x10<sup>3</sup> | 2.4x10<sup>-3</sup> ± 4.2x10<sup>-5</sup> | 0.94 |

Values reported represent the average of the global fit ( ± error) to the data from at least three independent BLI kinetic assays performed with different N protein preparations. Kinetics parameters were determined by using the bivalent analyte (1:2) binding model.

The robust and consistent expression of SARS-CoV-2 and HCoV-OC43 N on the surface of infected and surrounding cells suggests a conserved evolutionary function. As previously described for SARS-CoV-2 N (*21*), HCoV-OC43 N blocked CXCL12β-induced migration in chemotaxis assays with monocyte-like cells (MonoMac-1) in a concentration-dependent manner (Fig. 5B). N did not inhibit migration by itself, suggesting active blocking of CXCL12β-mediated migration. Although the calculated affinity of HCoV-OC43 N for CXCL12β (K_D_ = 1.5×10^-6^ ± 3.8×10^-8^ M) is 10-fold lower than SARS-CoV-2 N (K_D_ = 1.7×10^-7^ ± 1.2×10^-8^ M) (*24, 25, 34, 35*), it was only slightly less efficient on a molar basis at blocking chemotaxis.

SARS-CoV-1 and MERS-CoV N proteins also inhibit CXCL12β-mediated migration of MonoMac-1 cells (*21*). Endemic HCoV N proteins have a low amino acid sequence homology with HCoV-OC43 N (NL63 N, 42.7%; 229E N, 29.6%; HKU1 N, 63.8%). Regardless, these three HCoV N proteins blocked CXCL12β-induced migration in chemotaxis assays (Fig. 5C). Despite having the highest sequence homology with HCoV-OC43 N, HKU1 N showed lower inhibitory capacity than its Betacoronavirus counterpart. None of the endemic HCoV N proteins inhibited cell migration in the absence of CXCL12β, consistent with actively blocking CXCL12β.

Together, these findings indicate that N from both pathogenic and seasonal HCoVs block *in vitro* CHK-mediated leukocyte migration, consistent with the hypothesis that secreted N blocks CHK to facilitate initial viral replication and ultimately, host transmission.

### Surface HCoV-OC43 N is a target for Fc-mediated immunity during infection

Anti-N Abs binding to surface SARS-CoV-2 N activates Fc receptor-bearing cells and can reduce viral replication *in vivo* (*21, 36–38*). To analyze the anti-viral potential of polyclonal anti-N Abs acting via Ab dependent cellular cytotoxicity (ADCC), we generated Abs by immunizing mice with recombinant HCoV-OC43 N and tested them on HCoV-OC43 infected cells using FcγRIIIa receptor-expressing Jurkat reporter cells. HEK-293FT, BHK-21, MRC-5 and Vero infected cells each significantly activated Jurkat reporter cells in the presence of N-immunized pooled mice sera compared to infected cells incubated with pooled sera from naïve mice (Fig. 5D).

These results show that HCoV-OC43 N is a likely target for ADCC *in vivo* that potentially contribute to viral clearance and recovery.

## DISCUSSION

Here, we show that HCoV-OC43 N is stably bound to the surface of both N producing and neighboring cells, and that this is an intrinsic property of biosynthesized N as it robustly occurs in cells expressing N from a transgene. Levels of surface N on infected cells equal or exceed surface S in some cell types, in part, due to the retention of a variable fraction of S in the early secretory pathway, a common characteristic of HCoVs’ life cycle (*39*). HCoV-OC43 infected HAE cells expressed significant, but lower amounts of surface N compared to SARS-CoV-2 infected HAE, likely related to its limited replication in HAE cells (*40, 41*).

Since HCoVs N lack an endoplasmic reticulum targeting sequence and are not glycosylated or detected in the secretory pathway, N is almost certainly exported by a non-canonical pathway (*42*), like other cell surface viral nucleic acid binding proteins (e.g., HIV-Tat) (*43, 44*). N binding to the cell surface is specifically mediated through electrostatic interactions with HS/H by N RNA-binding domains. Several cellular proteins non-canonically secreted to the cell surface (e.g., FGF2, tau) also bind HS, which has been implicated in their membrane translocation (*45, 46*). The N signal detected 24 h post HCoV-OC43 infection of Vero cells was greater than on BHK-21 cells, which correlates with a higher amount of surface HS/H, supporting a role for HS/H in N export.

The reported affinity of HCoV-OC43 N protein for RNA is K_D_ = 1.2×10^-8^ M (*34, 35*). Here we show that HCoV-OC43 N affinity for HS, K_D_ = 1.5×10^-8^ M, is similar to its affinity for RNA, but much higher for H, K_D_ = 8.9×10^-10^ M. This is likely due to the higher sulfation of H compared to HS, thus its higher negative charge (*26*). Although SARS-CoV-2 N affinity for RNA,K_D_ = 8.3×10^-6^ M, (*24, 25*), is significantly lower than HCoV-OC43 N-RNA, we also reported a much higher affinity for HS/H (*21*). The high affinity of HCoV N proteins for H/HS may be key for their robust cell surface expression.

HCoV are not alone in exploiting HS/H as critical factors for viral attachment, entry, and immune modulation (*47, 48*). HCoVs interact with HS through the S protein, such as HCoV-NL63 (*49, 50*), and SARS-CoV-2 (*51, 52*). N is typically the most abundantly expressed HCoV protein (*53*), and its transfer to neighboring non-infected cells may amplify its contributions to viral fitness. Other secreted viral proteins also bind to the cell surface of infected or neighboring cells through HS/H (e.g., herpes simplex virus 2 glycoprotein gG (*54*), the myxoma virus T1 protein (*55*), the ectromelia virus E163 protein (*56*), the vaccinia virus B18 protein (*57*), and the molluscum contagiosum virus MC54L protein (*58*)).

Like N, CHKs are immobilized on secretory infected or neighboring cells by binding to HS/H. As a measure of their importance in anti-viral immunity, a number of viruses have evolved CHK-binding proteins to inhibit CHK activity, typically through interaction with the GAG- or receptor-binding domain of CHKs (e.g., myxoma virus M-T7, vaccinia virus A41, ectromelia virus E163, murine gammaherpesvirus-68 M3 and animal alphaherpesviruses gG) (*47, 48*). How the interaction between HCoVs N and CHKs affects chemotaxis *in vivo* remains to be determined. It is plausible that sequestration of CHKs by N limits their ability to recruit innate immune cells by limiting their capacity to diffuse from the site of infection, or by blocking their interaction with innate immune cells present at infection sites. Equally plausible, N-based display of biologically active CHKs may somehow exploit the local innate immune response to ultimately favor HCoV transmission between hosts. The effects of N on the wide array of CHKs bound need not be identical. Indeed, the different affinity of N from assorted HCoVs for the various CHKs may reflect selective effects on biological activity tailored to the special characteristics of each virus.

We show that as for SARS-CoV-2 N (*21, 36, 38*), HCoV-OC43 N is a target for Fc-based ADCC. Anti-N Abs reduced replication and protected mice from mouse hepatitis coronavirus disease and COVID-19 (*22, 36, 37, 59*). Since Abs, and T cells, to N cross-react among HCoVs, it is likely that prior infections with any HCoV provide at least some N-based protection to subsequent infection with a different HCoV. Indeed, the absence of HCoV-OC43 anti-N Abs is associated with severe COVID-19 (*60*). On other hand, presence of IgG Abs to alphacoronaviruses N proteins correlates with more severe COVID19 (*61*). Further, in patients with severe COVID-19, antibody responses biased toward the N at the expense of S, is associated with severe COVID-19 (*62–64*). Understanding the basis for these outcomes will be no easy matter.

Still, the strong immunogenicity of N and its antigenic stability makes it an attractive target for next-generation multivalent HCoV vaccines. A recent challenge study in rodents found that adding N mRNA to the standard S mRNA vaccine broadened protection against SARS-CoV-2 Delta and Omicron (*59*), though the bulk of the additive effect appears to be due to an anti-N CD8+ T cell response.

In summary, our findings support conserved roles for N as a target for Ab-mediated effector function in HCoV, and as a manipulator of the immediate CHK anti-viral response to localized infection.

## MATERIALS AND METHODS

### Cells

Vero cells (# CCL-81), BHK-21 (# CCL-10), CHO-K1 (# CCL-61), CHO-pgsA-745 (# CRL-2242), CHO-pgsB-618 (# CRL-2241), CHO-pgsE-606 (# CRL-2246), HEK293-FT (# CRL-11268), Rhabdomyosarcoma (RD) cells (# CCL-136), and MRC-5 (# CCL-171) cells were from the American Type Culture Collection (ATCC). MonoMac-1 cells (# ACC 252) were from the DSMZ-German Collection of Microorganisms and Cell Cultures. Vero, BHK-21, MRC-5 and HEK293-FT cells were grown in DMEM with GlutaMAX (Thermo Fisher # 10566016). CHO-K1, CHO-pgsA-745, CHO-pgsB-618, and CHO-pgsE-606 cells were grown in F-12K medium (Thermo Fisher # 21127022). MonoMac-1 cells were grown in RPMI 1640 (Thermo Fisher # 11875119). All cell media were supplemented with 8% (v/v) not heat inactivated FBS (Hyclone # SH30071.03). Cells were cultured at 37° C with 5% CO_2_. Cells were passaged at ~80-90% confluence and seeded as detailed for each individual assays. MucilAir™ (Epithelix # EP02MP) and SmallAir™ (Epithelix # EP21SA) HAE reconstituted from human primary cells obtained from nasal or bronchial biopsies were maintained in air-liquid interphase with specific culture medium (Epithelix # EP04MM, # EP64SA), in Costar Transwell inserts (Corning, NY, USA) per the manufacturer’s instructions.

### Virus preparation

HCoV-OC43 (# VR-1558) was obtained from ATCC. SARS-CoV-2 (isolate USA-WA1/2020, # NR-52281) was obtained from BEI resources. HCoV-OC43 was propagated in MRC-5 or RD cells at 35° C. SARS-CoV-2 was propagated by the NIAID SARS-CoV-2 Virology Core Laboratory under BSL-3 conditions using Vero (CCL-81) or Vero cells overexpressing human TMPRSS2 cells at 37° C. Both viruses were cultured in DMEM supplemented with GlutaMAX, 2% FBS, penicillin, streptomycin, and fungizone. Virus stocks were sequenced and subjected to minor variant analysis to ensure their sequence fidelity. The median tissue culture infectious dose (TCID_50_) and plaque forming units (PFU)/ml of viruses in clarified culture medium was determined on Vero cells after staining with crystal violet. SARS-CoV-2 infections were performed in the NIAID SARS-CoV-2 Virology Core BSL3 laboratory strictly adhering to its standard operative procedures.

### Abs and immunizations

For HCoV-OC43 N protein detection, we used sheep anti-OC43 N polyclonal Ab (MRC Protein Phosphorylation and Ubiquitylation Unit # DA116), and mouse anti-OC43 N monoclonal Ab (Sigma-Aldrich # MAB9013). Mouse immunization to obtain polyclonal anti-OC43 S or N sera were performed as followed: groups of five 8-to-12-week C57B6 mice (Taconic Farms Inc) were immunized with 4 μg/mouse of HCoV-OC43 S-His (Sino Biological # 40607-V08B) or N-His protein (Sino Biological # 40643-V07E) diluted in DPBS, adjuvanted by TiterMax® Gold (MilliporeSigma # T2684) (2:1) in 50 μl volume via intramuscular injections. Serum was collected 21 d after booster immunization, incubated at 56° C for 30 min, aliquoted, and stored at 4° C. Sera from each group were pooled, titrated, and their specificity was tested by flow cytometry (Fig. S9). Donkey anti-sheep IgG Alexa Fluor 488-conjugated (Thermo # A-11015), 647-conjugated (Thermo # # A-21448), goat anti-mouse IgG Alexa Fluor 488-conjugated (Thermo # A-11001) and 647 (# A-21235) were used as a secondary Abs.

### Plasmids

Plasmid pLVX-EF1alpha-eGFP-2xStrep-IRES-Puro encoding eGFP was obtained from Addgene (# 141395). The codon-optimized HCoV-OC43 N sequence was amplified from plasmid # 151960 (Addgene) with primers HCoV-OC43-N c-opt(pLVX)_Fw (gaattcgccgccaccatgtccttcaccccggg) and HCoV-OC43-N c-opt(pLVX)_Rv (cccgccgccttcgaggatctccgaagtgtcctcgg). The backbone vector pLVX-EF1alpha-2xStrep-IRES-Puro was linearized by PCR from Addgene plasmid # 141395 with primers pLVX-EF1alpha_Fw (ctcgaaggcggcggg) and pLVX-EF1alpha_Rv (ggtggcggcgaattc). Then, the HCoV-OC43 N sequence was cloned into backbone vector pLVX-EF1alpha-2xStrep-IRES-Puro by In-Fusion cloning (Takara Bio, Inc.), obtaining pLVX-EF1alpha-OC43-N-2xStrep-IRES-Puro.

### Immunofluorescence

For confocal microscopy imaging, 2.5 x 10^4^ cells were seeded on 12 mm glass coverslips in 24-well plates in the indicated medium supplemented with gentamycin (25 μg/ml) overnight. Cells were infected with HCoV-OC43 at a multiplicity of infection (MOI) of 1 PFU/cell for 1 h at 35° C. Virus was aspirated, and cells incubated in regular growth media. After 24 h, cells were washed with DPBS (Gibco # 14190-144) containing 5% goat serum (Jackson ImmunoResearch Labs. 005-000-121). Primary and secondary Abs were incubated with live cells at 4° C for 30 min. Cells were then washed twice with DPBS/5% goat serum and fixed in 4% PFA DPBS for 30 min at room temp. After fixation, coverslips were washed with DPBS and finally with deionized _i_H_2_O, and mounted with Dapi Fluoromount G™ mounting medium (VWR # 102092-102). Images were acquired with a Leica STELLARIS 8 confocal microscope platform equipped with ultraviolet and white light lasers, using a 63x oil immersion objective (Leica Microsystems # 11513859), with a 1x zoom resolution of 512 x 512 pixels. Maximum intensity projections were processed from z-stacks (at least 15 0.3 μm z-steps per image); and for background correction (Gaussian filter) and color processing, using Imaris (Bitplane). Background levels of signal for each cell type were set based on mock-infected stained conditions. Mock-infected coverslips were processed in parallel with infected counterparts, and infected coverslips were also incubated with all secondary-Abs-only as controls, and images were acquired using identical photomultiplier and laser settings.

### Flow cytometry

For cell surface protein expression analyses, 1 x 10^5^ cells were seeded on 24-well plates and mock or infected at an MOI of 1 PFU/cell for 1 h at 35° C, followed by aspirating the virus inoculum and adding medium containing 2% FBS. To infect HAE, the apical surface was washed thrice with DPBS and infected with 150 μl of viral inoculum (MOI = 1 for 3 h at 35° C). After the indicated hpi, cells were washed with DPBS, disaggregated with TrypLE™ Express Enzyme (Thermo Fisher # 12604039) for 5 min at 37° C, transferred to a 96 well plate and washed with HBSS (Lonza # 10-527F) with 0.1% BSA. HAE were dissociated according to the manufacturer’s instructions. In brief, the apical surface was washed thrice with DPBS and transferred to a new 24-well plate containing 1 ml of TrypLE™ Express. 250 μl of TrypLE™ Express were added to the apical surface for 15 min at 37° C, transferred to a 96 well plate and washed with HBSS 0.1% BSA. Cells were stained live with primary and secondary Abs and LIVE/DEAD™ Fixable Violet Dead Cell Stain Kit (Thermo Fisher # L34964) in DPBS, for 25 min at 4° C. After Ab staining, cells were twice washed with HBSS 0.1% BSA and analyzed. SARS-CoV-2-infected cells were fixed in 4% PFA DPBS for 30 min at room temp. PFA was aspirated, and cells resuspended in HBSS 0.1% BSA for analysis.

For transient surface protein expression, 2 x 10^5^ cells were seeded in 12-well plates and transiently transfected with 2 μg of a plasmid encoding HCoV-OC43 N or eGFP with TransIT-LT1 Transfection Reagent (Mirus Bio). At indicated time post transfection, cells were processed as described above for cell surface protein binding assays.

For cell surface protein binding assays using recombinant proteins, indicated cells were disaggregated, washed with DPBS, and 1 x 10^5^ cells were transferred to 96-well plates. Indicated amounts of recombinant GFP-His (Thermo Fisher # A42613) or HCoV-OC43 N-His (Sino Biological # 40643-V07E) were resuspended in 100 μl of DPBS, and cells were incubated for 15 min at 37° C and orbital shaking of 150 rpm. Cells were washed twice and stained as described above for subsequent flow cytometry analysis.

For electric charge neutralization assays, after infection (MOI = 10 for MRC-5 cells, MOI = 1 for CHO cells) or incubation with recombinant proteins, cells were washed twice, and incubated with 10 μg/ml of polybrene (MilliporeSigma # TR-1003-G) in DPBS for 15 min at 37° C and orbital shaking of 150 rpm. Cells were then washed twice and stained as described above for subsequent analysis.

For heparinases assays, 1 x 10^5^ cells in 96-well plates were treated with Bacteroides heparinase I (4.8 units), II (1.6 units) and III (0.28 units) (NEB # P0735S, # P0736S, # P0737S) in DPBS for 1 h at 30° C. Cells were washed twice, incubated with recombinant proteins, washed, and stained as described above for subsequent analysis.

For every assay and condition, at least 30,000 cells (100,000 cells for HAE) were analyzed on an BD FACSCelesta™ Cell Analyzer (BD Biosciences) with a high-throughput system unit, and quadrants in double staining plots were set based on mock-infected condition for each cell type. Data were analyzed with FlowJo (Tree Star) and plotted with Prism v9.1.1 software (GraphPad).

### N protein transfer assays

For transfected and non-transfected cells co-culture assays, 1 x 10^5^ cells were seeded on 6-well plates and transiently transfected with 2 μg of plasmids encoding HCoV-OC43 N or eGFP with TransIT-LT1. After 24 h, 9 x 10^5^ non-transfected cells were stained with CellTrace™ Violet and co-seeded with transfected cells and co-cultured for 12 h. Cells were then washed with DPBS and stained live *in situ* with primary and secondary Abs and LIVE/DEAD™ Fixable Violet Dead Cell Stain Kit, in DPBS for 25 min at 4° C. Cells were washed twice with HBSS 0.1% BSA, disaggregated with TrypLE™ Express Enzyme for 5 min at 37° C, transferred to 96 wells, washed, and resuspended in HBSS 0.1% BSA for flow cytometry analysis. For every assay and condition, at least 100,000 cells were analyzed on an BD FACSCelesta™ Cell Analyzer with a high-throughput system unit, and data were analyzed with FlowJo and plotted with GraphPad Prism software.

### CKs and GAGs

Recombinant human CKs used in this study (CCL1, CCL2, CCL3, CCL3L1, CCL4, CCL4L1, CCL5, CCL7, CCL8, CCL11, CCL13, CCL14, CCL15, CCL16, CCL17, CCL18, CCL19, CCL20, CCL21, CCL22, CCL23, CCL24, CCL25, CCL26, CCL27, CCL28, CXCL1, CXCL2, CXCL3, CXCL4, CXCL5, CXCL6, CXCL7, CXCL8, CXCL9, CXCL10, CXCL11, CXCL12α, CXCL12β, CXCL13, CXCL14, CXCL16, XCL1, CX3CL1, IL-1α, IL-1β, IL-6, IL-6Rα, IL-10, IL-12p70, IL-13, IL-17a, IL-18BP-Fc, IL-23, IL-27, IL-35, TNF-α, TNF-β, IFN-β, IFN-γ, IFN-λ1, IFN-ω) from PeproTech, and (IFN-α2 and IL-18) from Sino Biological, were reconstituted in DPBS 0.1% BSA at 10 μM, aliquoted and stored at −80° C. Heparin (# 2106), heparan sulfate from bovine kidney (# H7640), chondroitin sulphate A (# C9819) and chondroitin sulphate B (# C3788) were obtained from MilliporeSigma. Heparan sulfate from porcine mucosa (# AMS.GAG-HS01) and keratan sulfate (# AMS.CSR-NAKPS2-SHC-1) were purchased from ASMBIO. We assumed an average molecular weight of 30 kDa for heparan sulfate from porcine mucosa and 15 kDa for heparin (*65*).

### BLI assays

High-throughput-screening (HTS) binding assays were performed on an Octet Red384 (ForteBio) instrument at 30° C with shaking at 1,000 rpm. Streptavidin (SA) biosensors (Sartorius # 18-5019) were hydrated for 15 min in kinetics buffer (DPBS, 1% BSA, 0.05% Tween-20). GFP and HCoV-OC43 N proteins (2X-StrepTag tagged) in lysis buffer from crude lysates of transfected cells (see details below) were loaded into SA biosensors up to 5 nm of binding response for 300-600 s, prior to baseline equilibration for 180 s in kinetics buffer. Association of each analyte in kinetics buffer at indicated concentration was carried out for 300 s, followed by dissociation for 300 s or longer. Standard binding and kinetic assays between HCoV-OC43 N and GAGs or CKs were performed as described above for binding assays. Negative signal of N binding to GAGs, expected given the large size of heparin molecules, was flipped prior further analysis (*33, 66*). The data were baseline subtracted prior to fitting performed using the homogeneous (1:1) and heterogeneous binding models (2:1, mass transport, 1:2) within the ForteBio Data Analysis HT software v12.0.1.55. Mean K_D_, k_on_, k_off_ values were determined with a global fit applied to all data. The performance of each binding model fitting to the data was assessed based on the lowest sum of the squared deviations or measure of error between the experimental data and the fitted line (χ^2^), and the highest correlation between fit and experimental data (R^2^).

Experiments were repeated with at least three independently produced batches of recombinant protein in crude lysates, obtained from 30 x 10^6^ HEK293-FT cells transfected with 30 μg of plasmids encoding eGFP or HCoV-OC43 N protein (2X-StrepTag tagged) with TransIT-LT1. After 24 h, transfected cells were selected with 10 μg/ml of puromycin (Invivogen # ant-pr-1). After 48 h, transfected cells were disaggregated with TrypLE™ Express, washed with DPBS and lysated for 30 min at 4° C in 1 ml of lysis buffer (50 mM Tris-HCl pH 7.4, 150 mM NaCl, 5 mM KCl, 5 mM MgCl_2_, 1% NP-40 and 1x protease inhibitors (Roche # 4693159001), followed by centrifugation at 1000 x *g* at 4° C. Clarified supernatants (crude lysates) were collected, aliquoted, stored at - 20° C and characterized by immunoblotting (Fig. S8B), using IRDye® 680RD Streptavidin (LI-COR # 926-68079). HCoV-OC43 rN was additionally characterized by using mouse anti-OC43 N mAb (Sigma-Aldrich # MAB9013) followed by IRDye® 800CW Goat anti-Mouse IgG Secondary Ab (LI-COR # 926-32210).

### Chemotaxis assays

Recombinant human CXCL12β (3 nM), alone or in combination with purified recombinant proteins were placed in the lower chamber of a 96-well ChemoTx System plate (Neuro Probe # 101-5) in RPMI 1640 1 % FBS. As internal controls within each assay, medium or recombinant protein alone were used. MonoMac-1 cells (1.25 x 10^5^) were placed on the upper compartment and separated from the lower chamber by a 5 μm pore size filter. The cells were incubated at 37° C for 3 h in a humidified incubator with 5% CO_2_. The migrated cells in the lower chamber were stained with 5 μl of CellTiter 96 AQueous One Solution Cell Proliferation Assay (Promega # G3580) for 2.5 h at 37° C with 5% CO_2_, measuring absorbance at 490 nm using a Synergy H1 plate reader (Bio-Tek). The following recombinant proteins were used: SARS-CoV-2 N-His (Sino Biological # 40588-V07E), HCoV-OC43 N-His (Sino Biological # 40643-V07E), GFP-His (Thermo Fisher # A42613), NL63 N-His (Sino Biological # 40641-V07E), 229E N-His (Sino Biological # 40640-V07E), and HKU1 N-His (Sino Biological # 40642-V07E).

### ADCC reporter assay

For each indicated cell type, 2.5 x 10^4^ cells were seeded on 96-well flat white tissue culture-treated plates (Thermo Fisher # 136101), cultured overnight, and infected with HCoV-OC43 at an MOI of 1 PFU/cell (target cells) at 35° C. At 24 hpi, infected target cells were washed with DPBS and the medium was replaced with 40 μl of RPMI 1640 with 4% low IgG serum (Promega # G711A) containing 5 x 10^4^ Jurkat effector cells (Promega # G701A) and 10 μl of pooled mouse sera from naïve or immunized mice with HCoV-OC43 N-His. After overnight incubation at 37° C with 5% CO_2_, 50 μl of Bright-Glo™ Luciferase Assay lysis/substrate buffer (Promega # E2620) were added and luminescence was measured after 10 min using a Synergy H1 plate reader (Bio-Tek) with the following parameters: gain 150; measurement interval time 0.1 s; and integration time 0.01 s. Measurements were performed in triplicate and relative luciferase units (RLU) were plotted and analyzed with GraphPad Prism software.

### Statistical analysis

Statistical analyses were performed using GraphPad Prism software. When indicated, *p* values were calculated using Student ‘s two-tailed unpaired *t* test (at 95% confidence interval) and *p* < 0.05 was considered statistically significant. One-way ANOVA and Dunnett ‘s Multiple comparison test (at 95% confidence interval) was used to compare all conditions against untreated or mock-infected cells (as indicated for each case), considering *p* < 0.05 as statistically significant.

## Supporting information

Supplementary material Lopez-Munoz et al 2023

## ACKNOWLEDGMENTS

We thank James S. Gibbs for outstanding scientific and technical assistance. We are grateful to the NIAID SARS-CoV-2 Virology Core BSL3 Laboratory staff (Reed Johnson and Nicole Lackemeyer) for their training, help, and support.

## FUNDING

This work was supported by the Division of Intramural Research of the National Institute of Allergy and Infectious Diseases.

## AUTHOR CONTRIBUTIONS

Conceptualization: ADLM, JWY

Methodology: ADLM, JJSS

Investigation: ADLM

Formal analysis: ADLM

Visualization: ADLM

Resources: ADLM, JJSS

Supervision: JWY

Writing—original draft: ADLM

Writing—review & editing: ADLM, JJSS, JWY

## COMPETING INTERESTS

The authors declare no competing interests.

## DATA AND MATERIALS AVAILABILITY

All data needed to evaluate the conclusions in the paper are present in the paper and/or the Supplementary Materials.

## Notes

### Competing Interest Statement

The authors have declared no competing interest.

