## Supplementary material Lopez-Munoz et al 2023 for "Cell Surface Nucleocapsid Protein Expression: A Betacoronavirus Immunomodulatory Strategy"

**This PDF file includes:**

Figs. S1 to S9

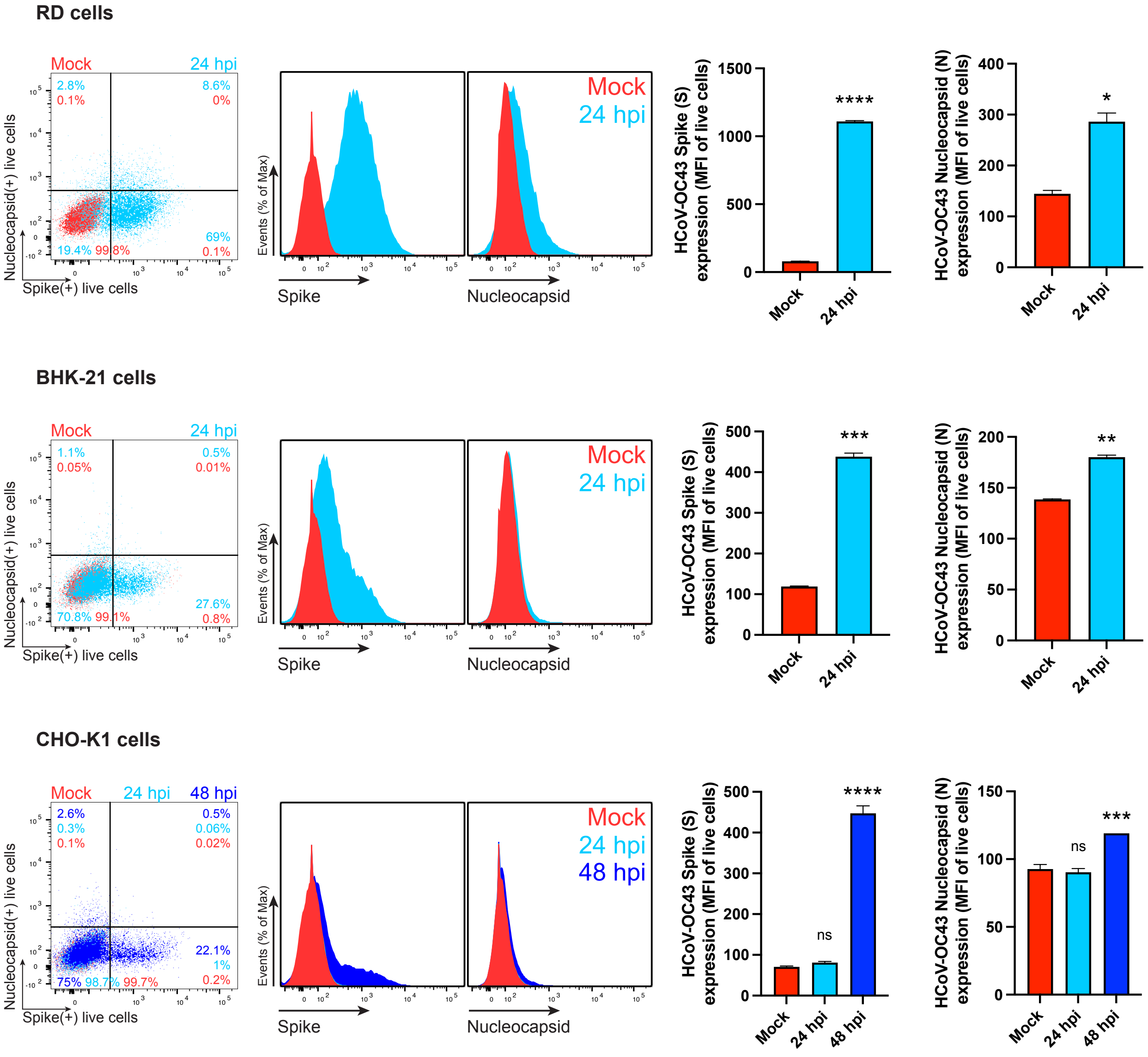


**Fig. S1. HCoV-OC43 N is present on the cell surface of additional cell types.** Flow cytometry analyses of RD, BHK-21 and CHO-K1 cells inoculated with HCoV-OC43 (MOI = 1), stained live with Abs at 24 hpi against the S and N proteins. For each cell type and infection, the following is shown: representative dot plots of flow cytometry analyses showing double staining of surface S and N, indicating the percentage of the gated cell population for each quadrant; as well as histogram overlays of surface S and N proteins. The MFI is plotted showing mean +/- SEM (n = 3). Student s two-tailed unpaired *t*-test was used to compare infected versus mock-infected RD and BHK-21 cells: ns (nonsignificant) $p > 0.05$, * $p < 0.05$, ** $p < 0.01$, *** $p < 0.001$, **** $p < 0.0001$. One-way ANOVA and Dunnett s Multiple comparison test were used for CHO-K1 cells: ns (nonsignificant) $p > 0.05$, *** $p < 0.001$, **** $p < 0.0001$. Data are representative of one experiment out of at least three independent experiments, each performed in triplicate.

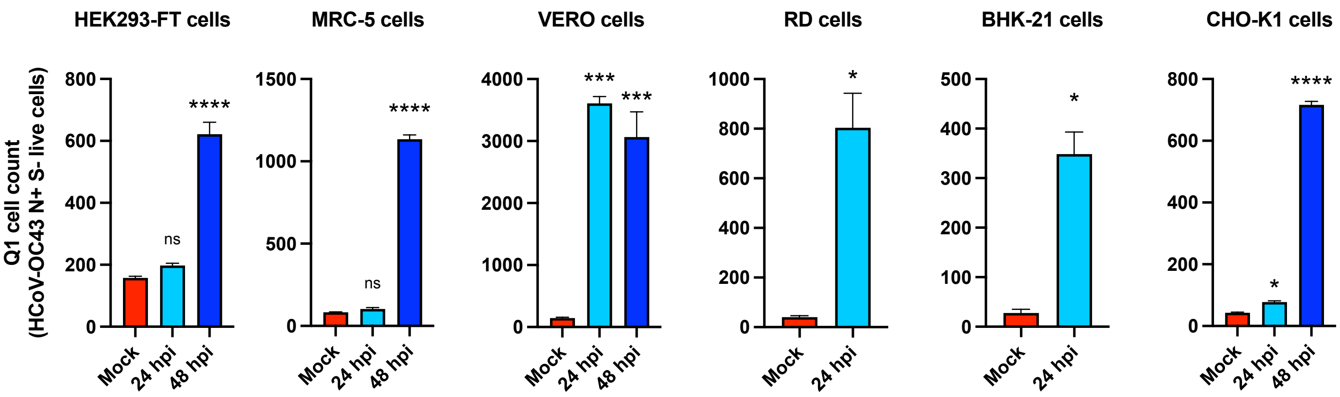


**Fig. S2. Additional evidence supporting N transfer from donor to recipient neighboring cells.** The number of cells expressing N but not S increased during infection in a time-dependent manner. Quadrant 1 (Q1) in dot plots of flow cytometry analyses showing double staining of surface N and S/eGFP identifies a cell subset of cells only expressing N during infection (Fig. 1, S1). Bar histograms show mean +/- SEM (n = 3) from Q1 cell counts of cells at 24 and 48 hpi HCoV-OC43 (MOI = 1). Data are representative of one experiment out of at least three independent experiments performed in triplicate.

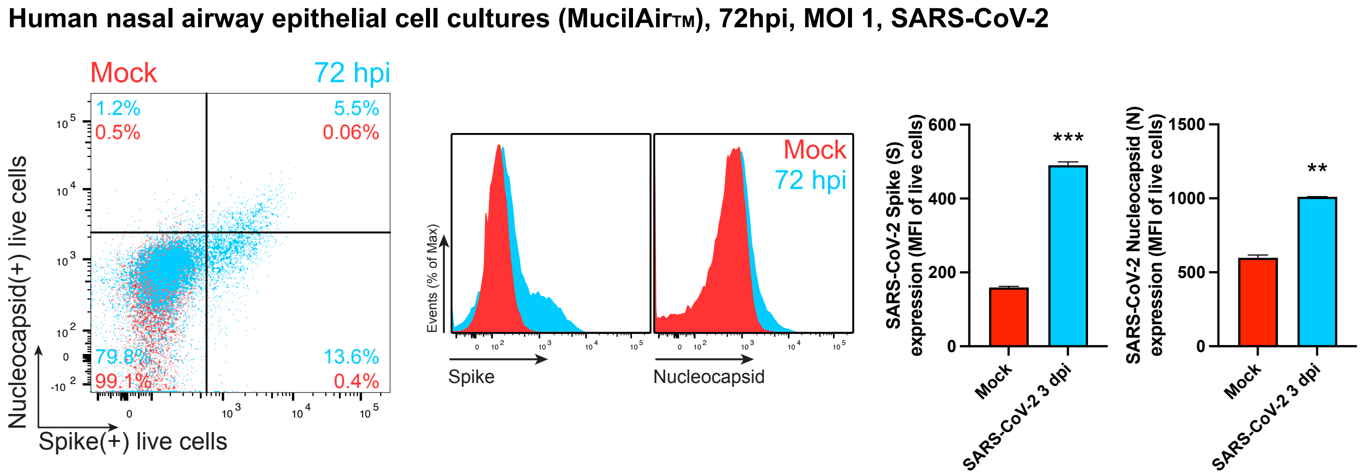


**Fig. S3. SARS-CoV-2 N is expressed on the surface of live infected HAE cells.** Flow cytometry analyses of human nasal airway epithelial cells infected with SARS-CoV-2 (MOI = 1), stained live at 72 hpi against the S and N proteins. Representative dot plot of flow cytometry analyses showing double staining of surface S and N, indicating the percentage of the gated cell population for each quadrant of the double staining. The following is shown: histogram overlays of surface S and N proteins, as well as the MFI is plotted showing mean +/- SEM (n = 2). ** $p < 0.01$, *** $p < 0.001$ by Student´s two-tailed unpaired *t*-test. Data are representative of one experiment out of two independent experiments performed in duplicate.

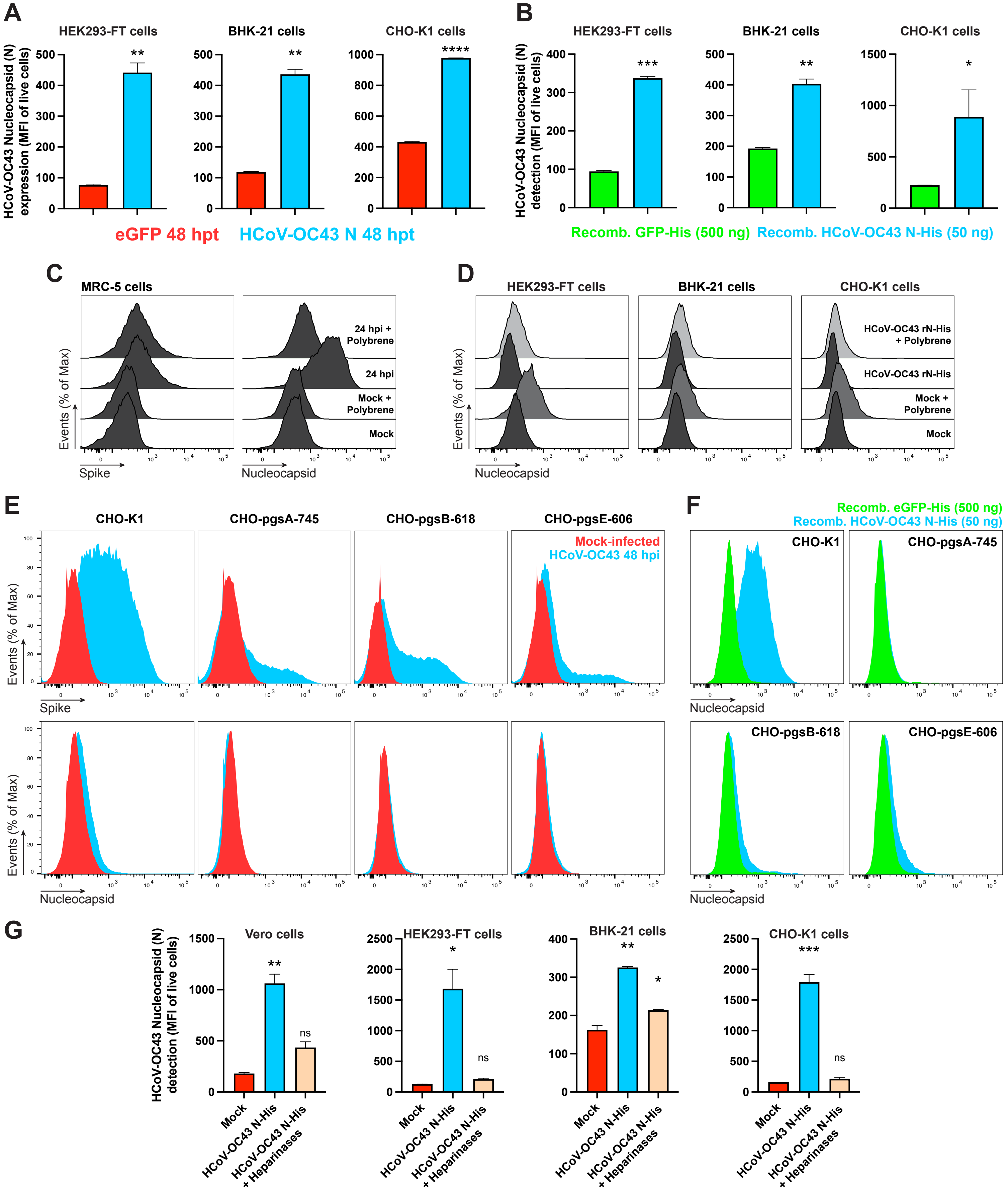


**Fig. S4. Other HCoV-OC43 genes are not required for N cell surface expression and binding to surface HS/H. (A)** Flow cytometry analyses of surface N expression in live HEK293-FT, BHK-21 and CHO-K1 cells transiently transfected with a plasmid encoding eGFP or N protein, detected with Abs. **(B)** Flow cytometry analyses of exogenous rN binding to HEK293-FT, BHK-21 and CHO-K1 cells, incubated with recombinant eGFP or N protein (50 ng) for 15 min, washed twice, and stained live with Abs. **(C)** Flow cytometry histogram semi-overlays of MRC-5 cells infected with HCoV-OC43 (MOI = 10), washed twice, incubated with 10 µg/ml of polybrene, washed twice again, stained live with Abs, and analyzed at 24 hpi. **(D)** Histogram semi-overlays of HEK293-FT, BHK-21 and CHO-K1 cells incubated with 50 ng of rN protein for 15 min, washed twice, incubated with 10 µg/ml of polybrene, washed twice, stained live with Abs, and analyzed by flow cytometry. **(E)** Histogram overlays of GAG-deficient CHO cells infected with HCoV-OC43 (MOI = 1), washed twice, incubated with 10 µg/ml of polybrene, washed twice again, stained live with Abs, and analyzed by flow at 48 hpi. **(F)** Histogram overlays of GAG-deficient CHO cells incubated with recombinant eGFP or rN protein for 15 min, washed twice, stained live with Abs, and analyzed by flow cytometry. **(G)** Flow cytometry analyses of Vero, HEK293-FT, BHK-21 and CHO-K1 cells treated with heparinases for 1 h, washed twice, incubated with 50 ng of rN protein for 15 min, washed twice again, stained live with Abs, and analyzed. The MFI of expressed **(A)** or bound **(B, G)** surface N protein from live cells is plotted for each case, showing mean +/- SEM (n = 2). In **(A, B)**, ns (nonsignificant statistically) $p > 0.05$, * $p < 0.05$, ** $p < 0.01$, *** $p < 0.001$, **** $p < 0.0001$ by Student s two-tailed unpaired *t*-test. In **(G)**, One-way ANOVA and Dunnett s Multiple comparison test were used to compare all conditions against untreated cells (mock): ns (nonsignificant) $p > 0.05$, * $p < 0.05$, ** $p < 0.01$, *** $p < 0.001$. All analyses were repeated with different protein preparations, and one representative assay out of at least three independent assays performed in duplicate is shown.

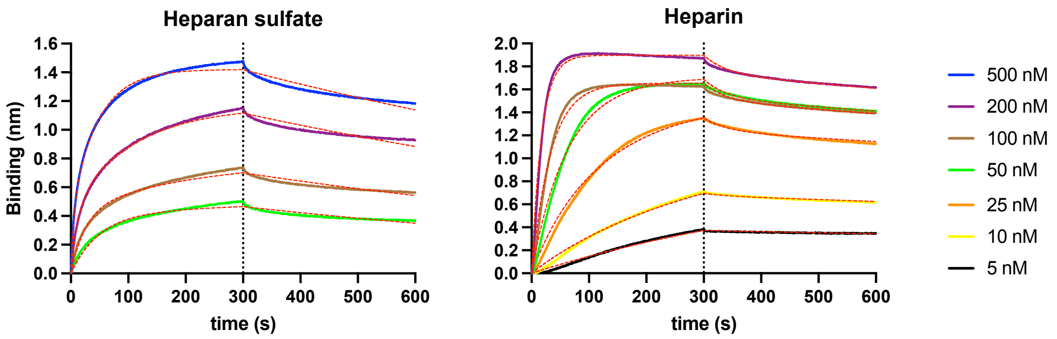


**Fig. S5.** BLI sensorgrams of kinetic assays depicting the interaction between immobilized HCoV-OC43 N protein and different concentrations of HS and H. All curves were analyzed with the ForteBio Data Analysis HT software, where red dashed lines correspond to a global fit of the data using the heterogeneous ligand (2:1) binding model. All analyses were repeated with different protein preparations, and one representative assay out of at least three independent assays performed in duplicate is shown.

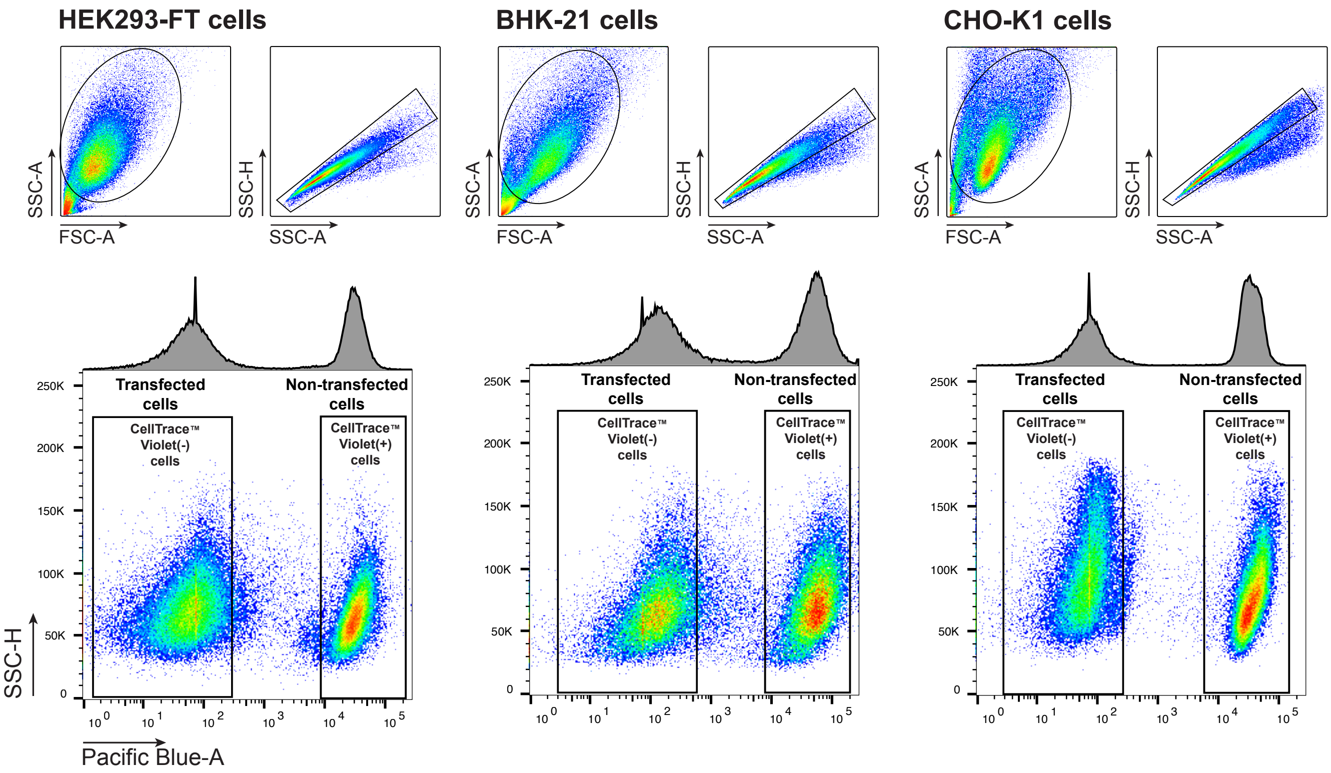


**Fig. S6. Gating strategy for HCoV-OC43 N protein transfer assays by transient transfection.** Gating and staining strategy for transiently transfected (donor) versus non-transfected (recipient) cell exclusion for flow cytometry analyses shown in **Fig. 4**. Non-transfected cells were stained with CellTraceTM Violet prior co-culture with transfected cells overnight. Representative dot plots of CellTraceTM Violet staining are shown, indicating the gating for transfected CellTraceTM Violet (-) cells, and non-transfected CellTraceTM Violet (+) cells. One representative experiment of at least three independent experiments performed in triplicate is shown.

A

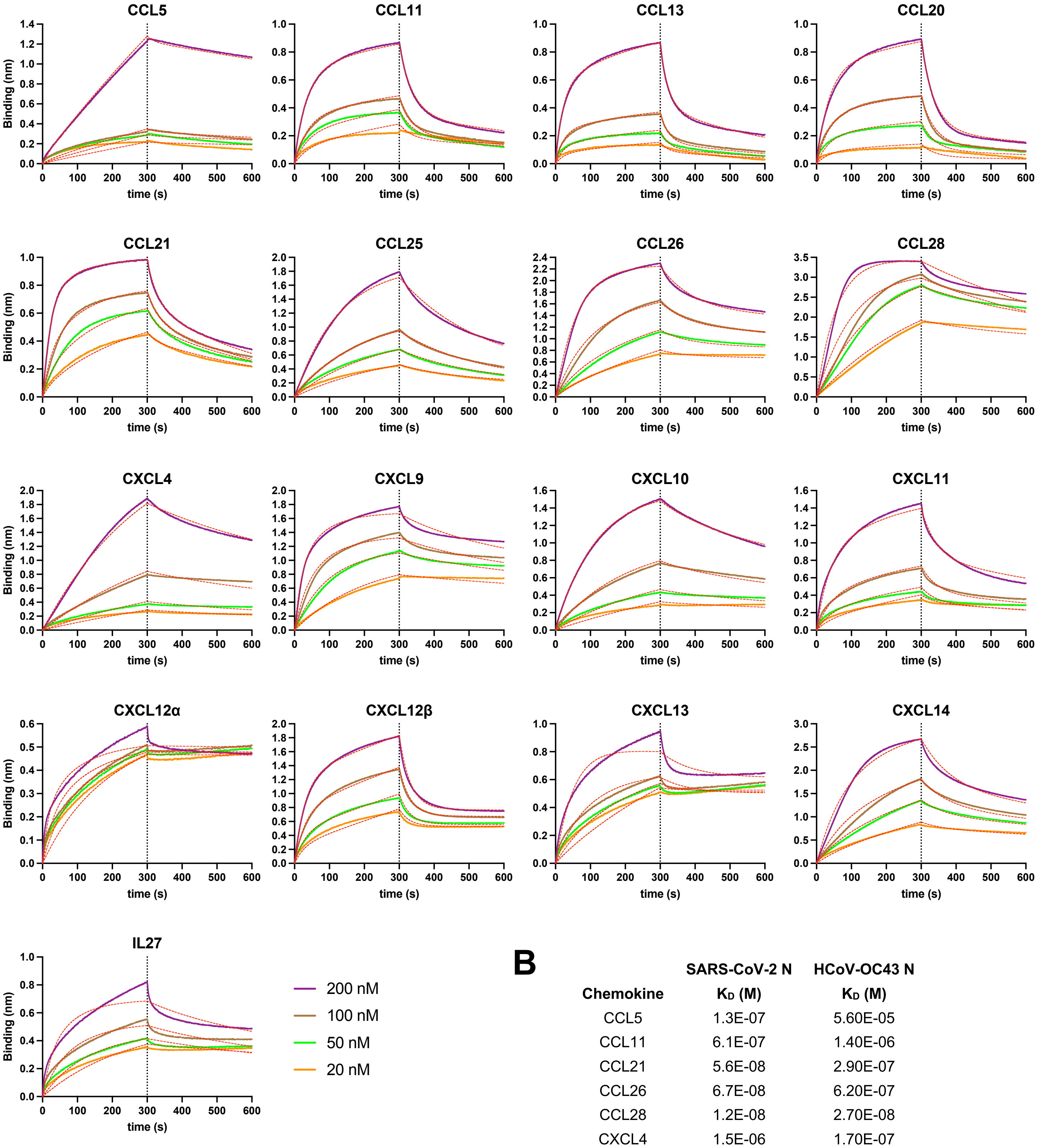


B

| Chemokine | SARS-CoV-2 N $K_D$ (M) | HCoV-OC43 N $K_D$ (M) |
|---|---|---|
| CCL5 | 1.3E-07 | 5.60E-05 |
| CCL11 | 6.1E-07 | 1.40E-06 |
| CCL21 | 5.6E-08 | 2.90E-07 |
| CCL26 | 6.7E-08 | 6.20E-07 |
| CCL28 | 1.2E-08 | 2.70E-08 |
| CXCL4 | 1.5E-06 | 1.70E-07 |
| CXCL9 | 3.3E-08 | 2.90E-08 |
| CXCL10 | 4.4E-08 | 3.70E-07 |
| CXCL11 | 1.6E-07 | 7.30E-07 |
| CXCL12B | 7.5E-07 | 1.50E-06 |
| CXCL14 | 1.1E-07 | 5.80E-07 |

**Fig. S7. HCoV-OC43 N protein specifically binds to 17 human CKs with high affinity. (A)** BLI sensorgrams of affinity kinetic assays between immobilized HCoV-OC43 rN protein and positively bound human CKs identified from BLI HTS binding assays. Sensorgrams show association and dissociation phases. The vertical dotted line indicates the end of the association step. Curves were analyzed with the ForteBio Data Analysis HT software, where red dashed lines correspond to a global fit of the data using the bivalent analyte binding model (1:2). All analyses were repeated with different protein preparations, and one representative assay out of at least three independent assays performed in duplicate is shown. **(B)** Comparative table showing kinetics affinities of HCoV-OC43 and SARS-CoV-2 N proteins for commonly bound CHKs.

**A**

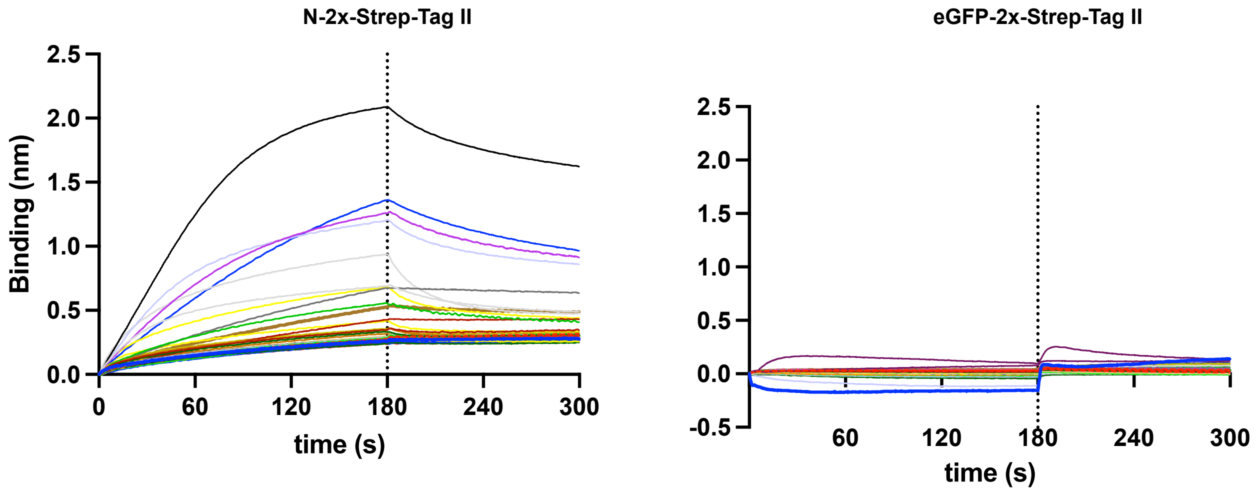


**B**

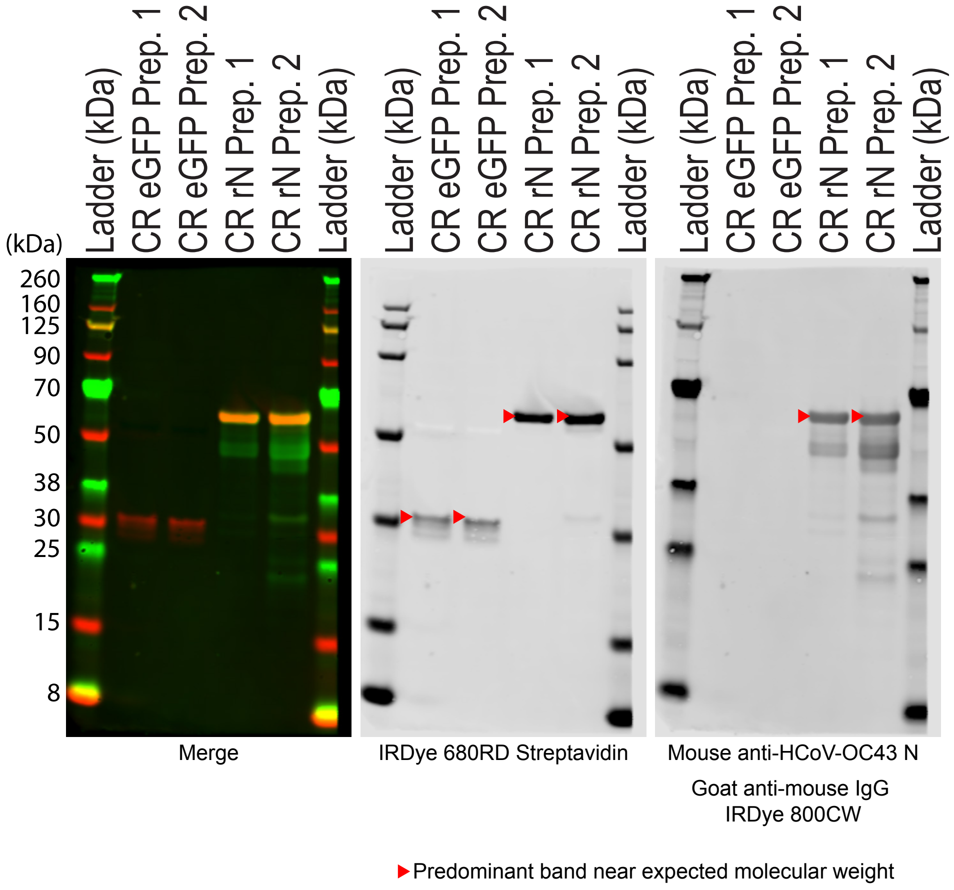


**Fig. S8. (A)** BLI sensorgrams of HTS binding assays between immobilized eGFP, and HCoV-OC43 N protein against 64 human cytokines at 100 nM (see detailed list in Material and Methods). N protein bound CCL5, CCL11, CCL13, CCL20, CCL21, CCL25, CCL26, CCL28, CXCL4, CXCL9, CXCL10, CXCL11, CXCL12α, CXCL12β, CXCL13, CXCL14 and IL27 across different independent assays. Sensorgrams show association and dissociation phases. The dotted line indicates the end of the association step. The analyses were repeated with different protein preparations and one representative assay of two independent HTS is shown. **(B)** Immunoblot detection of 2xStrep tag verified expression of predicted protein sizes (red arrowheads), which were loaded into streptavidin-coated biosensors for BLI HTS binding assays. Western blot images with molecular weight ladder from two different preparations of crude lysates (CR, 10 ul loaded) from transfected cells containing recombinant eGFP-2xStrep tag and HCoV-OC43 N-2xStrep tag. The same blot was stained with IRDye 680RD Streptavidin (LI-COR # 926-68079), and mouse anti-OC43 N mAb (Sigma-Aldrich # MAB9013) followed by IRDye® 800CW Goat anti-Mouse IgG Secondary Ab (LI-COR # 926-32210).

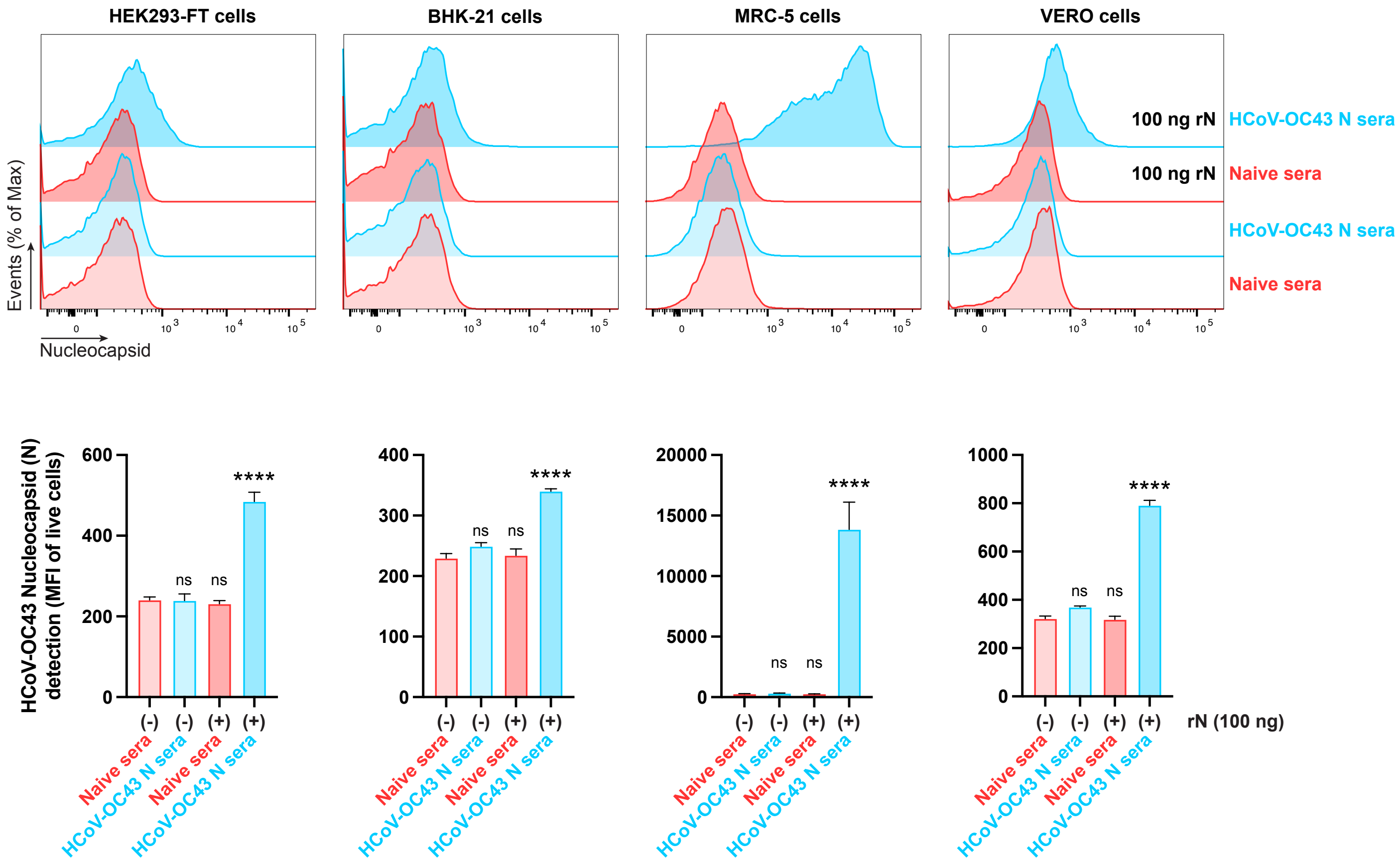


**Fig. S9. Characterization of pooled sera from naïve and immunized mice with HCoV-OC43 rN used for ADCC reporter bioassays.** Histogram semi-overlays of analyses of exogenous rN binding to HEK293-FT, BHK-21, MRC-5, and Vero cells, incubated with rN protein for 15 min, washed twice, stained live with pooled sera from naïve or immunized mice and secondary Abs, and analyzed by flow cytometry. The MFI is plotted showing mean +/- SEM (n = 3). One-way ANOVA and Dunnett s Multiple comparison test were used to compare all conditions against cells incubated only with sera from naïve mice: ns (nonsignificant) *p* > 0.05, **** *p* < 0.0001. Data are representative of one experiment out of two independent experiments performed in triplicate.
